# Selection shapes plant performance in a grassland biodiversity experiment

**DOI:** 10.1101/2024.08.19.608599

**Authors:** Francesca De Giorgi, Walter Durka, Yuanyuan Huang, Bernhard Schmid, Christiane Roscher

## Abstract

1. The increasing strength of positive biodiversity effects on plant community productivity in long-term biodiversity experiments has been shown to be related to mixed responses at species level. However, it is still not well understood if the varying environments in plant communities with different diversity also exert different selection pressures to which plant species respond adaptively or whether their responses are mainly due to phenotypic plasticity.
2. We conducted a transplant experiment for nine plant species in a 17-year-old grassland biodiversity experiment (Jena Experiment). We used offspring of plants *selected* in communities with different diversity or from plants which did not experience selection in the experiment (*naïve*). In a Community History Experiment, offspring of *selected* plants were planted in three test environments: their original plant communities with original soil, newly assembled plant communities with original soil, and newly assembled plant communities with new soil. Additionally, in a Selection Experiment we compared the performance of *selected* plants grown in their original environment to the performance of *naïve* plants grown in the same environment.
3. In all test environments, increasing species richness was on average associated with a decrease in plant individual biomass, reproductive output, relative growth rate, plant height, leaf greenness, and leaf nitrogen (N) concentration and an increase in SLA.
4. In the Community History Experiment survival was lower, while plant height, SLA, leaf N and C concentrations were highest in the test environment without plant and soil history. In high-diversity communities, individuals produced more biomass, grew taller and had higher leaf greenness in the original environment than in the other test environments.
5. In the Selection Experiment *selected* plants had weaker decline in their biomass, taller stature, and higher leaf C and N concentrations than
6. *naïve* plants with increasing species richness.
7. In both experiments, we found evidences of adaptive phenotypic responses. Specifically, in high-diversity communities, *selected* plants showed higher performance in their original environment, largely explained by plant community history and positive plant–soil feedbacks established over time.
8. *Synthesis:* Our study highlights the important role of eco-evolutionary feedbacks of species diversity, which affect plant performance over time.

## Introduction

Biodiversity is declining globally across different levels of organization; from genes to species to whole ecosystems (Cadotte et al., 2008; van der Plas, 2019). Concerns about the loss of biodiversity motivated the establishment of numerous biodiversity experiments. A major finding resulting from these experiments is that experimentally manipulated low-diversity communities show a decline in primary productivity and are less stable (Roscher et al., 2005). However, while increasing species diversity leads to an increased productivity at the community level, responses in biomass production of individual species are mixed and highly species-specific (Hille Ris Lambers et al., 2004; Schnitzer & Bongers, 2011; van Ruijven & Berendse, 2003). The environments experienced by species in plant communities of different diversity are characterized by different abiotic conditions and biotic interactions. Responses to these varying environments may be measurable in components of plant individual performance such as plant biomass, individual growth rate or reproductive output. For example, due to the greater neighbor biomass and density in species-rich communities, plants might invest more resources in vegetative growth and less in reproductive growth (Levins, 1968; Schmidtke et al., 2010). Moreover, previous studies found that species diversity negatively affected the proportion of flowering individuals, although the responses were highly species-specific (Lipowsky et al., 2011; Roeder et al., 2019). The taller and denser canopy in highly diverse communities typically induces responses to avoid shade, e.g. by increasing plant height, or to tolerate it, e.g. by increasing specific leaf area (SLA) (Bachmann et al., 2018; Lipowsky et al., 2011, 2015; Lorentzen et al., 2008).

Phenotypic responses of the same plant species growing either at high or low diversity may be a plastic adjustment or result of the selection of genotypes better fitting to their selection environment (Bailey et al., 2006; Linhart, 1988; Post & Palkovacs, 2009). Plasticity involves physiological, anatomical, or morphological changes within an organism’s lifetime that may improve its performance or survival in response to the original environment. In turn, adaptation occurs over the time of multiple generations and it requires a species to acquire or recombine genetic traits that improve performance or survival in its original environment (Demmig-Adams et al., 2008). Long-term biodiversity experiments allow us to study possible adaptation of plant populations to different original environments over the course of several generations.

Previous studies found that plants respond phenotypically to selection in high- vs. low-diversity communities, resulting in higher performance for plants selected in species-rich communities and the opposite for plants selected in species-poor ones. These findings suggest an important role of plant community diversity as a selective agent altering plant performance and trait expression (Dietrich et al., 2021; van Moorsel et al., 2018). Specifically, in grasslands more diverse communities are characterized by increased complementarity effects (Loreau & Hector, 2001), leading to a higher community productivity through the diversification of the use of space and light (Allan et al., 2011), partitioning of the soil resources (Fornara & Tilman, 2008; Roscher et al., 2008) and root depth distribution (Mueller et al., 2013). Another potential selective driver in plant communities of different diversity is the interaction of plant species with biotic and abiotic soil conditions, i.e. plant–soil feedback. For instance, monocultures generally experience a decline in productivity over time, attributable to a negative plant–soil feedback caused by the accumulation of host-specific pathogens (Schnitzer et al., 2011; Thakur et al., 2021). However, recent studies found that plants selected in monocultures can co- adapt to the associated soil community and benefit from mutualistic interactions (Zuppinger-Dingley et al., 2016). At high diversity positive plant– soil feedbacks are common (Eisenhauer et al., 2012; van Ruijven et al., 2020). The eco-evolutionary importance of plant–soil feedbacks can be disentangled thanks to the short generation time of both soil organisms and plants (Roeder et al., 2017; TerHorst & Zee, 2016). However, direct evidence on how plant diversity and soil organisms of original environments produce phenotypic responses of plant species, and to which extent they may be plastic adjustments or adaptations, is still lacking.

In this study, we used plots of the so-called ΔBEF Experiment established in a long-term grassland biodiversity experiment (Jena Experiment; Roscher et al., 2004; Vogel et al., 2019) to create test environments with different combinations of plant and soil community histories. Using these test environments, we aimed to disentangle the influences of plant diversity, as well as plant and soil community history on plant phenotypic responses. Taking advantage of the long-term experiment, we additionally aimed to separate the role of plasticity and adaptation in producing phenotypic responses. For this purpose, we established a transplant experiment to compare performance and trait expression of phytometers — test plants used to ”measure” their environment via survival, growth, and reproduction (Clements & Goldsmith, 1924; Strobl et al., 2018) — with different selection histories in the biodiversity experiment. Specifically, we studied the effects of selective pressures from two different perspectives. First, we focused on the influences of test environments with different community histories on the phenotypic responses of phytometers (Community History Experiment). One of the test environments, “with plant history, with soil history”, corresponded to the original environment in which the ancestors of phytometers had been selected for 17 years. The other two were test environments composed of newly assembled plant communities with soil used to establish the Jena Experiment (without plant history, with soil history) and newly assembled plant communities with new soil (without plant history, without soil history). Second, we focused on the effects of selection history on phenotypic responses of phytometers (Selection Experiment). For this purpose, we used again selected phytometers, whose ancestors underwent the selective pressures of the original environment “with plant history, with soil history”, and we transplanted them in the same original environment. At the same time, *naïve* phytometers, whose ancestors never experienced selective pressures in the biodiversity experiment, were transplanted in the same original environment “with plant history, with soil history”.

We hypothesized that:

1. The phytometers cultivated from plants that have undergone 17 years of selection in the biodiversity experiments show the highest performance when re-planted in their original environment (with plant history, with soil history). The performance is lower in the test environment “without plant history, with soil history” and the lowest in the test environment “without plant history, without soil history”. Responses in trait expression to increased plant diversity for these phytometers should be visible in all environments because of their selection history in communities with different diversity. This hypothesis is tested in the Community History Experiment.
2. Phytometers grown from *selected* plants have a higher performance in the original environment (with plant history, with soil history) than phytometers grown from *naïve* plants in the same environment. Furthermore, the responses of *selected* plants to increased plant diversity are more positive than the responses of *naïve* plants. This hypothesis is tested in the Selection Experiment.

## Materials and methods

### Experimental design of the Jena Experiment

The study was conducted in the Jena Experiment, a long-term biodiversity experiment located in the floodplain of the Saale River near Jena (Thuringia, Germany, 50°55′N, 11°35′E, 130 m a.s.l.) (Roscher et al., 2004). The experiment was established in 2002 on a former fertilized arable field. The soil on the site is a Eutric Fluvisol (FAO-Unesco 1997), whose texture varies from sandy loam to silty clay with increasing distance from the river. The experiment was organized in four blocks parallel to the riverside according to these soil characteristics (Roscher et al., 2004). A pool of 60 grassland species typically growing in meadows of the *Arrhenatherion* type (Ellenberg, 1988) was chosen for the experiment and classified into four functional groups: 16 grass species, 12 small herb species, 20 tall herb species and 12 legume species. The large species pool enabled the creation of different species mixtures, employing a near-orthogonal design to cross the experimental factors species richness (ranging from 1 to 60, including levels 1, 2, 4, 8, 16, and 60) and functional group number (1, 2, 3, or 4). In 82 plots of the main experiment, each species- richness level was represented with 16 replicates of different species compositions, with the exception of the 16-species mixtures which had only 14 replicates, and the 60-species mixture with one unique species composition but four replicates. Species for the different mixtures were randomly chosen from the respective functional groups. For the initial sowing of the experiment in May 2002, sowing density was 1000 viable seeds per m^2^ equally distributed among species in the mixtures. The seeds were purchased from a commercial supplier specialized in seeds of regional provenance (Rieger-Hofmann GmbH, Blaufelden-Raboldshausen, Germany). Some poorly established species were re-sown in autumn 2002 (for details see Roscher et al. 2004). Plots were mown twice per year (in early June and September) and the mown material was removed; weeding was generally carried out two to three times per year to maintain the intended species combinations. The experiment was not fertilized.

In this study, we used the ΔBEF Experiment (= Determinants of Long-Term Biodiversity Effects on Ecosystem Functioning) which was started in 2016 (Vogel et al., 2019). For the ΔBEF Experiment three subplots of 1.5 x 3 m size were established on each of the main experimental plots.

These subplots correspond to three test environments varying in soil history and plant community history. The first test environment, “with plant history, with soil history” (+PH+SH), was a subplot of the main experimental plot where communities were established in 2002. For the second test environment, the plant sod was removed with a digger, while the soil was mixed and homogenized to a depth of 30 cm on the respective subplot to obtain the combination “without plant history, with soil history” (-PH+SH). The third test environment, “without plant history, without soil history” (-PH-SH) was obtained by removing the vegetation and excavating the soil to 30 cm depth, which was replaced by soil from an adjacent arable field. For the second and third test environment, plastic sheets were installed as soil barriers in the top 30 cm of the soil to prevent a lateral mixing of the soils. Furthermore, for these two treatments plot-specific plant mixtures were sown in early May 2016, with a total density of 1000 viable seeds per m^-2^ equally divided among species using seeds purchased from the same supplier as for the initial establishment of the Jena Experiment in 2002.

### Study species

A subset of nine species representing the four functional groups of the Jena Experiment was chosen for the present phytometer experiments. The criteria for choice were a good representation of the species along the plant diversity gradient and the feasibility of collecting the required number of seed families from each community. The functional group “tall herbs” was represented by the species *Geranium pratense* L. (Geraniaceae), *Ranunculus acris* L. (Ranunculaceae) and *Crepis biennis* L. (Asteraceae), while “small herbs” were represented by *Plantago lanceolata* L. and *Plantago media* L. (both Plantaginaceae). *Lotus corniculatus* L. and *Medicago x varia* Martyn (both Fabaceae) were representing the “legumes”, and *Alopecurus pratensis* L. and *Trisetum flavescens* (L.) P. Beauv. (both Poaceae) were representing the “grasses”. *Crepis biennis* is monocarpic biennial to perennial, while the other eight species are perennials. To avoid the sampling of clonal replicates, we chose species whose individual genets were distinguishable for a long time after germination.

### Community History Experiment

The aim of the Community History Experiment was to compare the effects of different test environments, characterized by different community histories, on the performance of phytometers selected for 17 years in the same original environment (Fig. 1). For each phytometer species, 6 to 12 plots of different species richness, where the species belonged to the sown species combinations, were chosen. Our choice was limited by space, i.e. only one or two phytometer species could be planted per subplot of the ΔBEF experiment. However, for each species we aimed to include as many plots as possible, thus resulting in a dataset composed of nine species, distributed on a total of 54 plots. Out of the 54 plots, 23 of them hosted two species (Table S1). Seeds of the phytometer species were collected from four mother individuals in each plot (thus creating four seed families) in the original environment (+PH+SH) in 2019. The seeds were cleaned and stored at room temperature until the start of the phytometer experiments. In early January 2020, two to three seeds were placed in QuickPot^TM^ trays with 20 cm^3^ volume (Herrmann Meyer KG, Rellingen, Germany), which contained autoclaved (twice for 40 min at 121°C) soil from the field site mixed with sterile mineral sand (25 vol%). If more than one seedling germinated in a QuickPot, these were removed to allow the growth of single plantlets. To break dormancy and promote germination, seeds of *L. corniculatus* were scarified and seeds *R. acris* were treated with a solution of gibberellic acid (1000 mg L^-1^ for 24 hours) (Roscher et al., 2004). Afterwards, the pre-treated seeds of *R. acris* were germinated in petri dishes on moist filter paper, and the seedlings were transferred to the QuickPots when the radicle was visible. Pre-germination in petri dishes was also used for some seed families with low germination rates in other species to get enough seedlings. The phytometer plants were cultivated in a greenhouse (experimental field station Bad Lauchstädt, Saxony- Anhalt, Germany) at 18 °C during the day (14 hours) and 12 °C during the night (8 hours). In mid-March 2020, the trays were placed in a greenhouse with outside temperature and light conditions in order to harden the plants before being planted in the field. Between 4–15 April 2020 (see Table S1 for more details), three offspring per seed family were replanted into the same community of the original environment where they had been collected as seeds, which corresponded to the ΔBEF treatment “with plant history, with soil history” (+PH+SH). Three further offspring per seed family were transplanted into the test environments “without plant history, with soil history” (-PH+SH) and “without plant history, without soil history” (-PH-SH), respectively. In total, 36 plants (= 3 test environments x 4 seed families x 3 offspring) were planted per chosen plot and species. In a few cases, the number of seed families or offspring per seed family was not sufficient to get three offspring per subplot. In these cases the phytometer plants were distributed as equally as possible among the test environments. Planting distance was 20 cm with 25 cm distance from the plot margins; offspring of the seed families were randomly assigned to the planting positions. After planting, the phytometer plants were watered every second day for four weeks to assure a successful establishment. *Crepis biennis* phytometers were planted in the field in autumn (2 October 2020). The procedure and modalities followed the ones described before, with the germination and growth of the phytometers in the greenhouse starting in July 2020. The design resulted in a total of 2585 planted phytometers.

**Figure 1:**
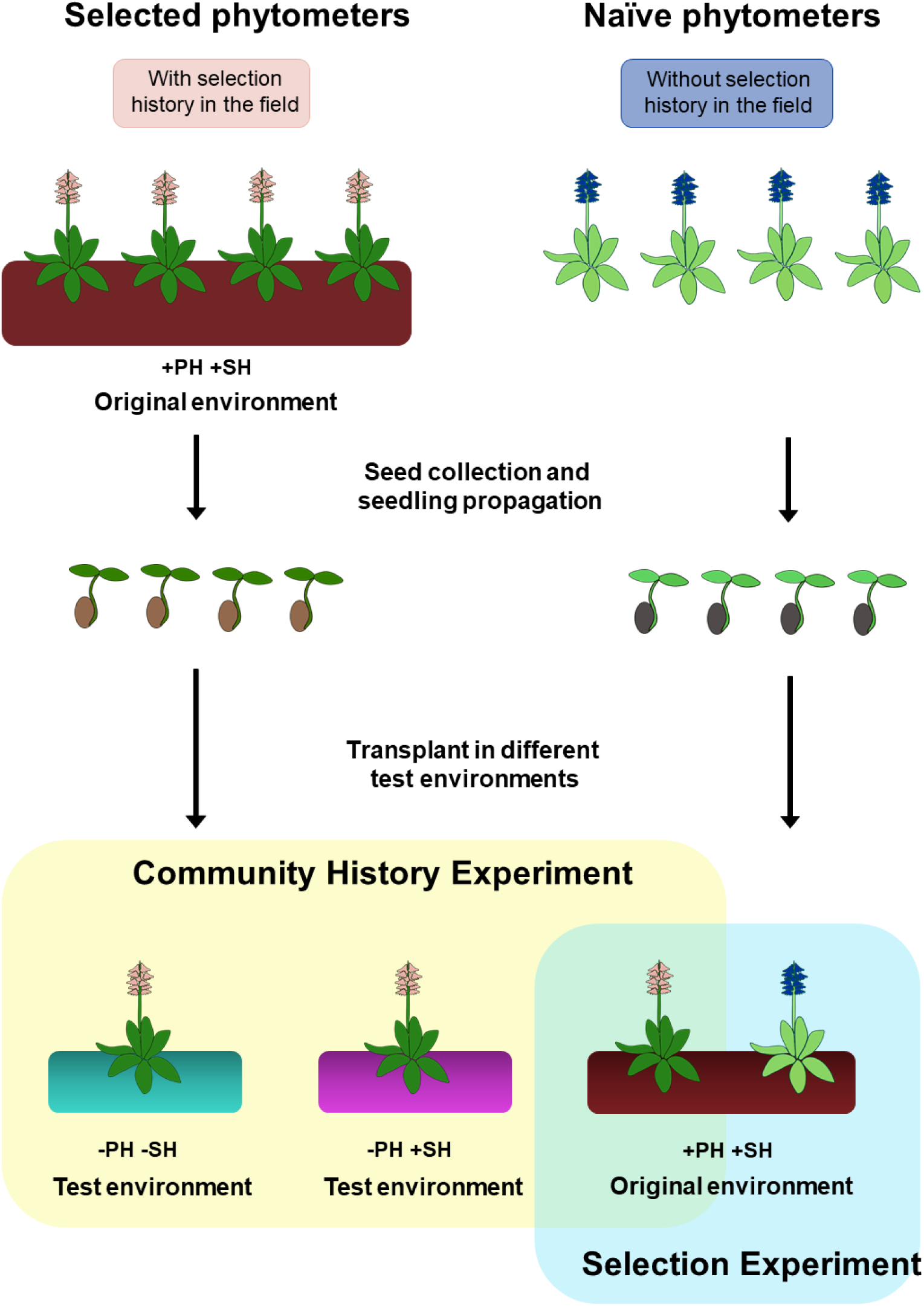
Design of the Community History Experiment and the Selection Experiment. Shown are the different phases of plant selection, seed germination and seedling propagation, and transplant in the field. Both experiments are shown at the species and plot level.

### Selection Experiment

The aim of the Selection Experiment was to compare the performance of phytometers whose ancestors experienced either the original environment in the biodiversity experiment (= *selected*) or not (= *naïve*) (Fig. 1). The selected phytometers were the same individuals used for the Community History Experiment in the original environment (+PH+SH), i.e. their original environments were the old communities sharing plant and soil history since 2002. The second group, composed by *naïve* phytometers, was obtained by growing plants from the initial seed lots used to establish the Jena Experiment in 2002. The ancestors of the *naïve* phytometers were germinated in summer 2018 from seeds that had been stored at –20°C since 2002. Seedlings were grown in a greenhouse with a standard substrate and planted outside in autumn in a sed bed at the Experimental Field Station Bad Lauchstädt. Seeds of these individuals were also collected in summer 2019. Seedlings of these seed families were grown and planted as *naïve* phytometers in the same original environment (+PH+SH) of the *selected* phytometers as described above for the Community History Experiment. This time, three offspring of the same four seed families per species (=12 phytometers per plot x species combination) were planted in all the plots, while seed families of the *selected* phytometers were plot-specific. In total, we planted 936 *naïve* phytometers, to be compared with 884 *selected* phytometers.

### Sampling and measurements

Measurements of phenotypic traits were taken simultaneously for both experiments. Before planting the phytometers in the field in April 2020, initial plant height was measured as the stretched shoot length or length of the longest leaf (in case of rosette plants). The number of leaves or shoots was counted, according to the morphology of the species (= time point t0). Between 18 August and 7 September 2020, about four months after planting in the field (= time point t1), we recorded the survival rate, considering a plant as alive when at least one green leaf could still be found. Then, stretched plant height was measured and the number of leaves were counted following the methods described above. Leaf greenness, an estimate of leaf chlorophyll concentrations was assessed by measuring the absorption of two different wave length (650 and 940 nm) with a portable chlorophyll meter (SPAD 502 Plus, Konica Minolta). Data were recorded for each individual by two averaged readings on different fully expanded leaves. Between 17–24 May 2021 (= time point t2), survival and plant height were recorded again. Afterwards, between 25 May and 10 June, one to five fully developed leaves per individual (depending on leaf and plant individual size) were sampled and stored in plastic bags in a cooling box until further processing. Leaf area was measured with a leaf area meter (LI-3000C Portable Leaf Area Meter equipped with LI3050C transparent belt conveyor accessory, LI-COR, USA) and samples were weighed after drying at 70 °C for 48 hours. Specific leaf area (SLA, mm ^2^ mg ^-1^) was calculated as the ratio between leaf area and dry mass. To determine nitrogen and carbon concentrations, leaves of the individuals composing a seed family for each subplot were pooled and milled to a fine powder with a ball mill (MM200, Retsch, Haan, Germany). After a second drying procedure for 24 h at 70 °C, sub-samples of the milled sample material were scanned with an MPA FT-NIR spectrometer (Bruker, Bremen, Germany) to determine nitrogen (mg N g_dw_ ^-1^) and carbon concentration (mg C g_dw_ ^-1^). To establish and validate the optimal NIRS model for predicting N (NLeaf) and C (CLeaf) concentrations, around 20 samples per species were processed by conventional chemical analysis using an elemental analyzer (Vario EL cube, Elementar Analysensysteme GmbH, Langenselbold, Germany). The NIRS calibration models used to predict N and C concentration in the samples had coefficients of determination r^2^ = 0.986 for N and r^2^ = 0.827 for C. The final trait measurements and biomass harvest took place between 18 August and 9 September 2021 (= time point t3), in order to have repeated measurements comparable across the years. First, survival was assessed and then the plant individuals were harvested at soil surface using a knife. Plants were stored in separate plastic bags in a cooling box until measurements took place in the laboratory. There, stretched plant height was measured, leaf or shoot number were counted and leaf greenness was recorded as above. Afterwards, plants were dried at 70 °C for 48 hours and weighed to get plant individual biomass. During both the field campaigns in spring and summer 2021, the presence of inflorescences for each individual was noted. Combining these two time points to account for the different species-specific flowering time during the growing season, we obtained an estimate of the individual reproductive status for the year 2021. Using the repeated measurements of leaf or shoot number taken during the first measurements in the field and the final harvest, we calculated relative growth rates for each individual as

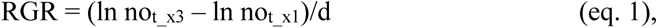

where, ln not_x3 and ln not_x1 represent the natural logarithms of the number of leaves or shoots counted at the end of the experiment in August/September 2021 (t3) and in August/September 2020 (t1), and d is the number of days between these time points (Harper, 1977). During all field campaigns, the height of the surrounding vegetation around the phytometers was measured in each subplot, as the average of three measurements across the area occupied by the experimental plants. For variables derived from the measurements see Table 1.

**Table 1:** Overview of the phenotypic traits measured on phytometer plants during the course of the experiment, where t0 = initial measurement at planting in spring 2020, t1 = summer 2020, t2 = spring 2021, and t3 = summer 2023 .

| Variable | Function | Unit | Description | Measured |  |  |  |
| --- | --- | --- | --- | --- | --- | --- | --- |
|  |  |  |  | t0 | t1 | t2 | t3 |
| Survival | Performance | / | Survival of the individual |  | x | x | x |
| Reproductive status | Performance | / | Presence or absence of flowers |  |  | x | x |
| Plant biomass | Performance | g | Total aboveground biomass |  |  |  | x |
| Leaf/Shoot number | Light acquisition | / | Number of shoots or leaves | x | x |  | x |
| Plant height | Light acquisition | cm | Length of the longest shoot or leaf | x | x | x | x |
| Leaf greenness | Light acquisition | / | Estimate of chlorophyll concentration |  | x |  | x |
| Specific leaf area | Light acquisition | mm<sup>2</sup><sub>leaf</sub> g<sup>-1</sup><sub>leaf</sub> | Leaf area per leaf dry mass |  |  | x |  |
| N<sub>Leaf</sub> | Nutrient acquisition | mg N g<sub>dw</sub><sup>-1</sup> | Leaf nitrogen concentration |  |  | x |  |
| C<sub>Leaf</sub> | Nutrient acquisition | mg C g<sub>dw</sub><sup>-1</sup> | Leaf carbon concentration |  |  | x |  |

### Data analyses

Data analysis was performed with R Statistical Software (v4.2.2; R Core Team 2022, http://www.R-project.org). To test our hypotheses, we performed analyses of variance. To test the effects of species richness and test environments with different community history on plant performance and trait expression for the Community History Experiment (H1), we optimized a model for all the response variables fitting different explanatory variables in the following order: block, sown species richness (log-linear), test environment of the ΔBEF experiment (factor with three levels) and its interaction with sown species richness, followed by the functional group identity of the phytometer species (factor with four levels) and its interaction with sown species richness and test environment, species identity of the phytometer species (factor with nine levels, when fitted after functional groups measuring variation within these among species) and its interaction with sown species richness and test environment, plot and its interaction with species identity, subplot and its interaction with species identity, and finally seed family identity and its interaction with test environment. Because of the nested experimental design, we calculated the F and P values using the appropriate error terms as in mixed models (for details, see Table S2). To investigate the role of test environment in determining the height of the surrounding vegetation of each plot, we created a model fitting block, sown species richness (log-linear), test environment and its interaction with sown species richness, followed by plot and subplot identity. Differences among the test environments and the four functional groups were identified using Tukey’s HSD test (R package *multcomp*) (Hothorn et al., 2008). To investigate the effects of species richness on trait expression and the effects of selection history on plant performance in the Selection Experiment (H2), we adjusted the previous model by substituting the term test environment as well as its interactions with a term for selection history, and by removing the term subplot together with its interactions, because the Selection Experiment only used data from the original environment (+PH+SH). Again, we calculated the F and P value with the appropriate error terms as in mixed models (Table S3).

For the analyses of binomial variables (i.e. survival and reproductive status) in both experiments we used generalized linear mixed-effects models with a binomial error distribution, implemented with the g*lmer* function in *lme4* package (Bates et al., 2015). We used type-I (sequential) sum of squares. As random terms composing the null model, we fitted plot, seed family, interaction between plot and species identity, subplot, interaction between subplot and species identity and interaction between test environment and seed family. Starting from the null model, fixed effects were added stepwise, including block, species richness (log-linear), test environment, and interaction between test environment and species richness. Finally, we created for the analyses of both experiments an alternative model in order to account for possible effects of light competition by the surrounding vegetation that could partly explain the effects of species richness. In these models, the canopy height of the surrounding vegetation was fitted as a covariate right after the block term.. When needed, the variables were log-transformed to meet the requirements of normality.

## Results

### Effects of species richness, functional groups, and seed families on plant performance and trait expression

In both the Community History Experiment and the Selection Experiment sown species richness affected plant performance and the expression of most measured traits. Specifically, in the Community History Experiment, plant performance measured as plant individual biomass, the proportion of plants reaching the reproductive stage, and relative growth rate (RGR) decreased with increasing species richness, while individual survival was not affected (Table 2, Table S5, Fig. 2A-C). Regarding traits related to light acquisition, plant height measured in the first year and leaf greenness measured in both years decreased, while SLA increased with species richness (Fig. 2 D, E, Fig. S1A, B). Plant height, measured in the second year, did not significantly respond to species richness. However, when we accounted for the canopy height of the surrounding vegetation before entering species richness into the model, species richness had a significant effect on plant height in 2021 (Table S4). For leaf elemental concentrations, we found that CLeaf showed little variation along the species-richness gradient, while NLeaf decreased with increasing species richness (Fig. 2F). Analyses of the Selection Experiment showed similar patterns of phenotypic responses to sown species richness (Table 3, Fig. S2).

**Figure 2:**
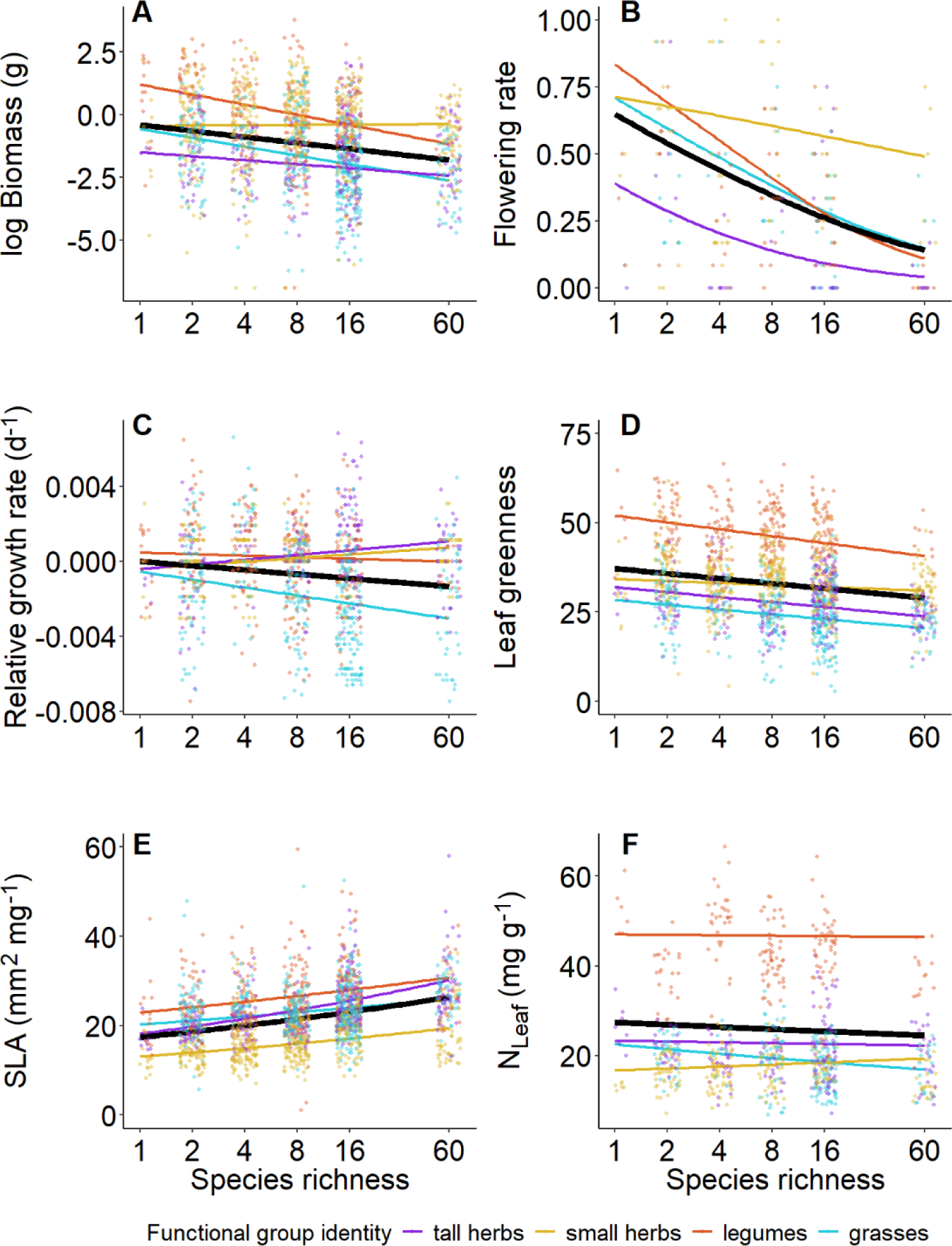
Effects of sown species richness on (A) plant individual biomass (summer 2021), (B) proportion of individuals reaching reproductive stage at the plot level (year 2021), (C) relative growth rate summer 2020 to summer 2021), (D) leaf greenness (summer 2021), (E) specific leaf area (SLA) (spring 2021), and (F) leaf nitrogen concentration (NLeaf) (spring 2021). Solid colored lines represent a significant relationship at the functional-group level, while the solid black line represents the mean response across all species.

**Table 2:** Results of linear models and generalized linear mixed effects models for the *Community History Experiment* testing effects of sown species richness (SR), treatment, their interaction, identity of the functional group (FG-ID), its interaction with species richness and treatment, species and its interaction with species richness and treatment on plant performance and trait expression. If variables were measured at different time points, it is indicated with t1 = summer 2020 and t3 = summer 2021. Shown are F and P values. Abbreviations of variable names: LeafG = leaf greenness, CLeaf = leaf carbon concentration, NLeaf = leaf nitrogen concentration, SLA = specific leaf area.

| Source of variation | Survival_t3 |  | Reproductive status |  | Biomass |  | RGR_t3-t1 |  |  |  |
| --- | --- | --- | --- | --- | --- | --- | --- | --- | --- | --- |
| | $\chi^2$ | P | $\chi^2$ | P | F | P | F | P | | |
| Block | 11.96 | 0.008 | 0.11 | 0.991 | 5.50 | 0.003 | 7.31 | 0.001 |  |  |
| Species richness (SR) | 0.48 | 0.490 | 10.57 | 0.001 | 25.76 | <0.001 | 7.31 | 0.010 |  |  |
| Treatment | 12.62 | 0.002 | 0.22 | 0.894 | 2.01 | 0.139 | 0.07 | 0.933 |  |  |
| SR x Treatment | 3.83 | 0.147 | 2.86 | 0.239 | 2.64 | 0.076 | 2.21 | 0.116 |  |  |
| FGID | 10.80 | 0.013 | 22.42 | <0.001 | 21.34 | <0.001 | 18.42 | <0.001 |  |  |
| SR x FGID | 13.87 | 0.003 | 2.29 | 0.515 | 5.61 | 0.023 | 7.83 | 0.025 |  |  |
| Treatment x FGID | 9.18 | 0.164 | 0.68 | 0.995 | 0.89 | 0.544 | 0.15 | 0.982 |  |  |
| Species-ID | 21.35 | 0.001 | 70.17 | <0.001 | 9.31 | 0.003 | 21.04 | 0.003 |  |  |
| SR x Species-ID | 4.31 | 0.506 | 14.01 | 0.016 | 4.55 | 0.029 | 1.58 | 0.304 |  |  |
| Treatment x Species-ID | 16.29 | 0.092 | 19.37 | 0.036 | 0.35 | 0.964 | 0.60 | 0.732 |  |  |
| Plot | / | / | / | / | 2.50 | <0.001 | 2.11 | 0.002 |  |  |
| Species-ID x Plot | / | / | / | / | 1.22 | 0.292 | 1.10 | 0.365 |  |  |
| Subplot | / | / | / | / | 2.27 | 0.007 | 2.21 | 0.029 |  |  |
| Species-ID x Subplot | / | / | / | / | 1.75 | 0.013 | 0.81 | 0.689 |  |  |
| SF | / | / | / | / | 1.66 | <0.001 | 0.90 | 0.761 |  |  |
| Treatment x SF | / | / | / | / | 0.66 | 1.000 | 1.08 | 0.237 |  |  |
| Source of variation | Plant height_t3 |  | LeafG_t3 |  | SLA |  | N<sub>Leaf</sub> |  | C<sub>Leaf</sub> |  |
|  | F | P | F | P | F | P | F | P | F | P |
| Block | 19.88 | <0.001 | 1.21 | 0.315 | 10.3 | <0.001 | 31.85 | <0.001 | 8.30 | <0.001 |
| Species richness (SR) | 1.01 | 0.319 | 45.51 | <0.001 | 62.6 | <0.001 | 12.25 | 0.001 | 0.00 | 1.000 |
| Treatment | 2.09 | 0.129 | 0.42 | 0.656 | 8.7 | <0.001 | 3.59 | 0.031 | 3.15 | 0.047 |
| SR x Treatment | 3.71 | 0.028 | 4.38 | 0.015 | 0.3 | 0.747 | 2.40 | 0.096 | 0.26 | 0.769 |
| FG-ID | 49.81 | <0.001 | 121.65 | <0.001 | 67.8 | <0.001 | 206.41 | <0.001 | 70.91 | <0.001 |
| SR x FG-ID | 8.92 | 0.006 | 1.00 | 0.441 | 1.2 | 0.356 | 2.67 | 0.119 | 2.00 | 0.193 |
| Treatment x FGID | 0.08 | 0.997 | 0.39 | 0.863 | 0.3 | 0.903 | 0.00 | 1.000 | 0.67 | 0.680 |
| Species-ID | 16.25 | 0.001 | 7.91 | 0.006 | 17.2 | <0.001 | 29.73 | <0.001 | 12.57 | 0.001 |
| SR x Species-ID | 1.57 | 0.272 | 1.27 | 0.361 | 1.1 | 0.415 | 1.33 | 0.341 | 1.03 | 0.461 |
| Treatment x Species-ID | 1.09 | 0.377 | 1.04 | 0.417 | 1.6 | 0.106 | 0.72 | 0.705 | 1.10 | 0.367 |
| Plot | 2.97 | <0.001 | 3.45 | <0.001 | 3.0 | <0.001 | 3.91 | <0.001 | 2.57 | <0.001 |
| Species-ID x Plot | 1.51 | 0.164 | 2.03 | 0.050 | 3.4 | 0.002 | 3.59 | 0.001 | 1.84 | 0.078 |
| Subplot | 1.92 | 0.026 | 0.96 | 0.570 | 1.4 | 0.164 | 1.06 | 0.444 | 0.62 | 0.956 |
| Species-ID x Subplot | 1.46 | 0.067 | 1.55 | 0.046 | 1.1 | 0.285 | 1.68 | 0.017 | 1.64 | 0.022 |
| SF | 1.07 | 0.285 | 0.99 | 0.518 | 1.0 | 0.420 |  | / | / | / |
| Treatment x SF | 0.96 | 0.677 | 1.16 | 0.056 | 0.9 | 0.729 |  | / | / | / |

Note: $\chi^2$ values of binomial variables (survival rate and reproductive status) were obtained using generalized mixed effects models. For other variables, F values were obtained performing an analysis of variance. Because of the nested design of the experiment, we calculated the F and P values using the appropriate error term, which can be found in Table S2.

**Table 3:** Results of linear models and generalized linear mixed effects models for the *Selection Experiment* testing effects of sown species richness (SR), selection history, functional group identity (FG-ID), species identity and their interactions with species richness and selection history on plant performance and trait expression. If variables were measured at different time points, it is indicated with t1 = summer 2020 and t3 = summer 2021. Shown are F and P values. Abbreviations of variable names: LeafG = leaf greenness, CLeaf = leaf carbon concentration, NLeaf = leaf nitrogen concentration, SLA = specific leaf area.

| Source of variation | Survival_t3 |  | Reproductive status |  | Biomass |  | RGR_t3-t1 |  |
| --- | --- | --- | --- | --- | --- | --- | --- | --- |
| | $\chi^2$ | P | $\chi^2$ | P | F | P | F | P |
| Block | 9.23 | 0.026 | 1.40 | 0.707 | 3.90 | 0.014 | 5.46 | 0.003 |
| Species richness (SR) | 1.49 | 0.223 | 6.64 | 0.010 | 9.57 | 0.003 | 1.58 | 0.215 |
| Selection | 1.12 | 0.289 | 0.00 | 0.956 | 28.33 | 0.003 | 4.20 | 0.133 |
| SR x Selection | 0.00 | 0.984 | 1.37 | 0.241 | 2.11 | 0.153 | 1.14 | 0.293 |
| FG-ID | 13.57 | 0.004 | 14.91 | 0.002 | 14.46 | <0.001 | 7.55 | <0.001 |
| SR x FG-ID | 7.50 | 0.058 | 5.92 | 0.116 | 5.08 | 0.035 | 2.08 | 0.245 |
| Selection x FG-ID | 0.25 | 0.969 | 3.87 | 0.276 | 0.82 | 0.521 | 0.42 | 0.751 |
| Species-ID | 21.61 | 0.001 | 57.82 | <0.001 | 10.79 | 0.003 | 6.28 | 0.054 |
| SR x Species-ID | 3.85 | 0.571 | 10.75 | 0.057 | 7.82 | 0.009 | 0.22 | 0.877 |
| Species x Selection | 19.92 | 0.001 | 15.31 | 0.009 | 0.42 | 0.836 | 0.44 | 0.727 |
| Plot | / | / | / | / | 3.87 | 0.033 | 1.26 | 0.463 |
| Species x Plot | / | / | / | / | 1.69 | 0.113 | 3.15 | 0.016 |
| Selection x Plot | / | / | / | / | 1.65 | 0.009 | 0.92 | 0.607 |
| Seed family | / | / | / | / | 1.09 | 0.215 | 1.17 | 0.106 |

| Source of variation | Plant height_t3 |  | Leaf greenness_t3 |  | Specific leaf area |  | N<sub>Leaf</sub> |  | C<sub>Leaf</sub> |  |
| --- | --- | --- | --- | --- | --- | --- | --- | --- | --- | --- |
|  | F | P | F | P | F | P | F | P | F | P |
| Block | 22.18 | <0.001 | 1.71 | 0.177 | 7.97 | <0.001 | 18.57 | <0.001 | 5.47 | 0.003 |
| Species richness (SR) | 0.00 | 1.000 | 24.18 | <0.001 | 48.29 | <0.001 | 19.47 | <0.001 | 4.21 | 0.046 |
| Selection | 6.25 | 0.054 | 3.41 | 0.124 | 0.18 | 0.689 | 0.83 | 0.403 | 3.33 | 0.127 |
| SR x Selection | 1.47 | 0.231 | 0.03 | 0.853 | 1.50 | 0.226 | 4.64 | 0.036 | 4.21 | 0.045 |
| FG-ID | 42.98 | <0.001 | 1145.75 | <0.001 | 54.83 | <0.001 | 143.20 | <0.001 | 44.02 | <0.001 |
| SR x FG-ID | 14.39 | 0.002 | 1.60 | 0.285 | 1.50 | 0.288 | 5.33 | 0.026 | 0.08 | 0.971 |
| Selection x FG-ID | 3.89 | 0.063 | 1.24 | 0.375 | 0.42 | 0.743 | 1.33 | 0.330 | 1.98 | 0.195 |
| Species-ID | 48.53 | <0.001 | 3.60 | 0.075 | 15.57 | 0.001 | 71.20 | <0.001 | 9.97 | 0.003 |
| SR x Species-ID | 6.30 | 0.016 | 1.14 | 0.431 | 1.26 | 0.368 | 2.80 | 0.094 | 5.39 | 0.018 |
| Species-ID x Selection | 1.35 | 0.245 | 1.29 | 0.271 | 1.41 | 0.223 | 3.07 | 0.010 | 0.42 | 0.838 |
| Plot | 5.61 | 0.011 | 1.57 | 0.302 | 0.96 | 0.581 | 2.98 | 0.052 | 2.17 | 0.122 |
| Species-ID x Plot | 1.45 | 0.189 | 1.90 | 0.083 | 4.32 | <0.001 | 1.28 | 0.254 | 0.76 | 0.640 |
| Plot x Selection | 1.72 | 0.005 | 1.12 | 0.297 | 1.09 | 0.333 | 1.10 | 0.301 | 1.03 | 0.424 |
| SF | 0.81 | 0.965 | 1.39 | 0.002 | 0.98 | 0.567 | / | / | / | / |

Note: $\chi^2$ values of binomial variables (survival rate and reproductive status) were obtained using generalized mixed effect models. For other variables, F values were obtained performing an analysis of variance. Because of the nested design of the experiment, we calculated the F and P values using the appropriate error term, which can be found in Table S2.

Phytometers belonging to different functional groups differed in all measured variables with the exception of survival (Table 2, 3, Table S8, S9). Moreover, species-richness effects showed some variation among functional groups (significant interactions species richness x functional group identity) in both experiments. In particular, for plant individual biomass, small herbs showed a positive response along the plant-diversity gradient, while for species of the other three groups plant individual biomass consistently declined with increasing species richness (Fig. 2A). Additionally, in spite of the general decrease in plant height with increasing species richness (in the Community History Experiment), both small and tall herbs increased in height (Fig. S1A). Functional group-specific responses were also detectable for leaf greenness measured in the first year, for which grasses were the only group with higher values of leaf greenness in high-diversity communities (Fig. S1B, Fig. S3A). In the Community History Experiment, RGR was positive for small and tall herbs (Fig. 2C). Finally, in contrast to the general decrease in NLeaf with increasing species richness, NLeaf of small herbs showed the opposite response in the Selection Experiment (Fig. S2E).

In both experiments, the nine phytometer species within functional groups differed in plant individual biomass and the expression of all traits. In contrast, the seed families used in the Community History Experiment, differed in plant individual biomass, but not in the expression of the measured traits. However, the seed families used for the Selection Experiment, comparing phytometers with and without selection history in the plots, differed in plant height in the first year and leaf greenness in the second year (Table 2). Note: χ2 values of binomial variables (survival rate and reproductive status) were obtained using generalized mixed effects models. For other variables, F values were obtained performing an analysis of variance. Because of the nested design of the experiment, we calculated the F and P values using the appropriate error term, which can be found in Table S2.

### Effects of soil and plant history on phytometer performance and trait expression

The ΔBEF test environment, in which the phytometers grew, affected their survival rates, height and expression of leaf traits (SLA, NLeaf, CLeaf) (Table 2, Fig. 3, Fig. S4). We found that, irrespective of species richness, survival rates significantly differed among test environments. The original environment +PH+SH did not have the highest survival rates, however, the difference was 3 to 6% from the test environment +PH+SH and 9 to 11 % from the test environment -PH+SH (Table S10, Fig. 3A). Plants grew taller, had larger SLA and higher NLeaf and CLeaf in the test environment -PH-SH (Table S10, Fig. 3D, E, Fig. S4 B). However, when accounting for variation in canopy height, only the treatment effects on plant height remained significant (Table S6). Additionally, we found a significant interaction between species richness and the test environments for plant height and leaf greenness in the second summer: in both cases, phytometers in the original environment +PH+SH were shorter and had lower leaf greenness in low-diversity communities, while those in high-diversity communities were taller and had higher leaf greenness when compared to the other two test environments (Table 2, Fig. 3B, Fig. S4B). When accounting for the effects of canopy height, the effects on plant individual biomass became significant, i.e. decrease in biomass in diverse communities was less pronounced in phytometers grown in their original environment +PH+SH (Table S6, Fig. S4C). Concerning the effects of the test environments on the height of the surrounding vegetation, we found that the -PH-SH had the highest canopy during spring 2021 compared to the other two test environments (Table S11, Fig. S5A, C, E).

**Figure 3:**
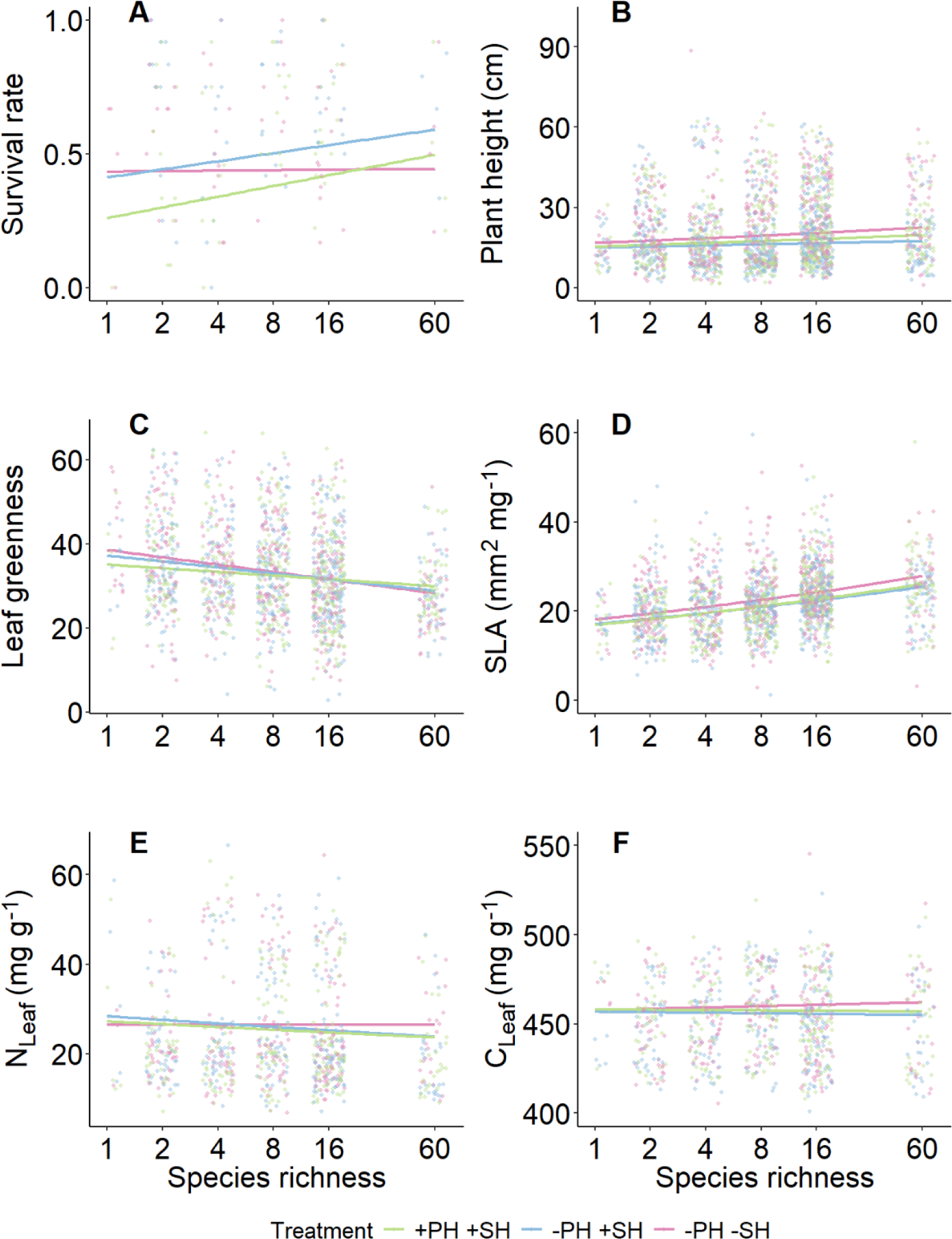
Effects of the test environments on (A) survival rate at the plot level (summer 2021), (B) plant height (spring 2021), (C) leaf greenness (summer 2021), (D) specific leaf area (spring 2021), (E) leaf nitrogen concentration (NLeaf) (spring 2021), and (F) leaf carbon concentration (CLeaf) (spring 2021). Solid colored lines represent a significant relationship at the treatment level. Shown are the different test environments of the ΔBEF Experiment. i.e. “with plant history, with soil history” (+PH+SH), “without plant history, with soil history” (-PH+SH) and “without plant history, without soil history” (-PH-SH)..

### Effects of selection history on phytometer performance and trait expression

*Selected* and *naïve* phytometers grown in the original environment +PH+SH differed in plant individual biomass and in plant height measured in the second year (Fig. 4). Specifically, *selected* phytometers grew taller, and they responded stronger to increased species richness by producing more biomass than the *naïve* ones. Moreover, *naïve* phytometers had higher NLeaf and CLeaf at low levels of species richness, while the *selected* ones had the highest nutrient concentrations at higher species richness (Table 3, Table S7, Fig. 4). In this experiment, the significant effect of selection history on plant individual biomass and plant height in the second year was mediated by canopy height of the surrounding vegetation (Table S6).

**Figure 4:**
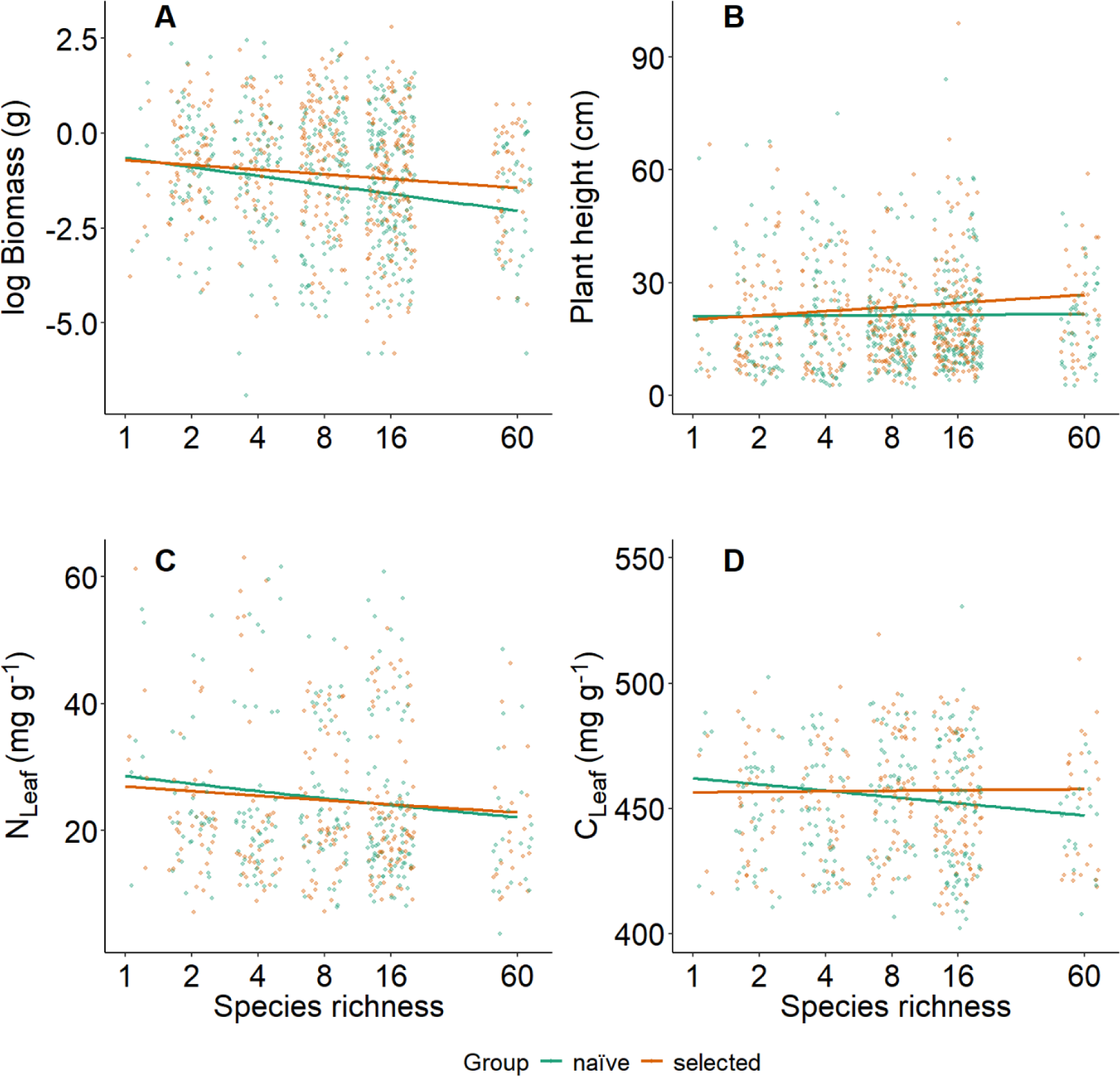
Effects of the selection history on (A) plant individual biomass (summer 2021), (B) plant height (summer 2021), (C) leaf carbon concentration (CLeaf) (spring 2021) and (D) nitrogen concentration (NLeaf) (spring 2021). Solid colored lines represent a significant relationship among the variables and selection history of the phytometers.

## Discussion

### Phenotypic responses to plant species diversity and differences among functional groups

In accordance with previous studies in the Jena Experiment (summarized in Roscher et al., 2018), we found that species richness significantly affected the expression of most of the studied traits. It is increasingly acknowledged that high plant species richness leads to a higher community-level productivity, largely explained by increased niche complementarity among species and positive plant–soil feedback (Barry et al., 2019; Thakur et al., 2021). However, performance of individual species is not always positively affected by plant diversity (Hille Ris Lambers et al., 2004; Hooper & Vitousek, 1997; Schnitzer et al., 2011; Thein et al., 2008; van Ruijven & Berendse, 2003). In our experiment we found that the rate of survival did not change with species richness, but on average individuals flowered less frequently, had a smaller RGR, and produced less biomass, suggesting a lower individual performance at higher plant diversity. However, these responses varied across species, as we found positive responses of small herbs to increasing species richness. We also found significant effects of species richness on trait expression such as lower height in the first year, increasing SLA, and decreasing leaf greenness and NLeaf. Confirming the role of light competition as an important driver of plant individual performance and trait expression (Bachmann et al., 2018; Thein et al., 2008), variation in plant height as well as the effects of species richness on plant height were largely explained by canopy height. Moreover, we observed that functional groups responded to some degree differently to increasing plant species richness. Specifically, small herbs produced more biomass, and their RGR and NLeaf were higher as a response to increasing species richness. Obviously, the small stature of this functional group does not allow plants to reach the upper canopy layers and acquire direct sun light, and their adjustment to canopy shade is crucial to increase performance. Morphological and physiological adjustment in trait expression may help them to cope with the highly variable supply of light during the different seasons (Roscher et al., 2011). The increase in RGR and NLeaf as well as leaves with larger SLA are often found in shaded plants (Valladares & Niinemets, 2008). The tall herbs, on their part, responded to the increased competition for light by increasing their height and RGR. This is line with several previous findings, which showed how species able to grow tall enough, invest their resources in increased height and biomass (Hirose & Werger, 1995). We conclude that the different growth responses to increased plant diversity, as found by the different functional groups and shaped by physiological and facilitation processes are essential for species coexistence, especially in highly productive and competitive communities.

### Modifications of phenotypic responses to plant species diversity in original environment vs. increasingly different test environments (Community History Experiment)

We tested the effects of different test environments, characterized by different plant and soil community history, on performance of phytometers selected for 17 years in the same original environment. We expected phytometers to perform better in their original environment +PH+SH “with plant history, with soil history”, and to have a lower performance in the test environments -PH+SH “without plant history, with soil history” and -PH-SH “without plant history, without soil history” (hypothesis H1). The test environment in which phytometers grew had indeed an influence on their survival rates as well as on the expression of several traits measured in the second growing season. Contrary to our expectations, survival was highest in the test environment -PH+SH. We found little evidence of an interaction of the test environments with species richness, expected based on previous findings, that showed how the presence of soil history is usually detrimental for plant performance in species-poor communities, mostly explained by the density-dependent accumulation of pathogens in the soil over time (Kulmatiski et al., 2012; Thakur et al., 2021). However, the interactions between interspecific competition and plant–soil feedbacks, especially across a resource gradient, can significantly alter plant performance (Lekberg et al., 2018; Zuppinger-Dingley et al., 2016). These effects depend on various factors and more field studies are needed to better disentangle the role of plant-soil history and their feedback for phenotypic responses in a community context. Concerning the measured traits, we found the largest effects in the test environment - PH-SH. As canopy height was the tallest in this test environment, phytometers were more likely shaded by taller plants than in the other two treatments. Likely, the greater height and SLA of the phytometers in this test environment could be a response to the shade caused by the surrounding vegetation. Several studies have shown that the presence of taller neighbors induces plants responses in order to either tolerate or avoid canopy shade, like increasing specific leaf area and plant height (Bachmann et al., 2018; Lipowsky et al., 2011, 2015; Lorentzen et al., 2008). Moreover, we found higher CLeaf and NLeaf in the test environment without plant and without soil history. The soil used to establish this treatment within the ΔBEF Experiment was taken from a neighboring agricultural field, in the same way as the soil of the field site of the Jena Experiment, the difference being the high level of fertilization and productivity (Vogel et al., 2019; Wagg et al., 2022). Thus, the test environment having new soil was more fertile than the test environments with soil history, leading to higher CLeaf and NLeaf.

We did not find a direct effect of species richness on plant height, but phytometers growing in their original environment increased their height with increasing species richness. We propose that the phytometers growing in their original environment were advantaged and consequently better competitors at high diversity compared with the phytometers growing in the other two test environments. In line with this result, we also found that, after accounting for canopy height, the reduction in phytometer biomass with increasing species richness was less pronounced in their original environment compared to the other two test environments. Positive effects of the original environment on plant performance were also found in previous studies (Dietrich et al., 2021; Johnson et al., 2010; Pregitzer et al., 2010; van Moorsel et al., 2018; Wagg et al., 2015). Based on all these findings, we suggest that history of selection in species-rich communities plays an important role in enhancing plant performance. In addition to the overall negative effects of increasing species richness on leaf greenness, we again identified a significant interaction between test environment and species richness. Values of leaf greenness were higher in the test environment -PH-SH, especially at low diversity. Given that soil fertility in the test environment without soil history was probably higher (see above), we found this advantage to be especially visible in low-diversity communities. This could likely be explained by lower interspecific competition and to the more sparse vegetation, which provided favorable conditions for phytometers in monocultures to effectively exploit nutrients.

In summary, the enhanced performance of phytometers in their original environment was clearly visible after accounting for potential competition for light, again highlighting the importance of light availability for plants adjustments to their environment, both trough plasticity and adaptation.

### Modification of phenotypic responses to plant species diversity in selected vs. naïve plants. (Selection Experiment)

Testing the effects of selection history on plant performance, we expected *selected* phytometers to perform better than the *naïve* ones in their original environment (hypothesis H2). We found some evidence supporting this hypothesis with regard to plant individual performance, measured as phytometer biomass and height. Irrespective of being *selected* or *naïve*, phytometers suffered from competition in highly diverse communities, resulting in smaller plants. However, this reduction in size was less pronounced in *selected* phytometers. Furthermore, we found increased CLeaf and NLeaf with increasing species richness in *selected* phytometers, and the opposite trend for the *naïve* ones. In summary, the performance and trait expression of the *naïve* phytometers were comparable (plant biomass and height) or better (CLeaf, NLeaf) than the *selected* ones in low-diversity communities, while they were worse in high-diversity communities.

Previous studies found that plants selected in monocultures can benefit from mutualistic interactions with the soil community over time, thus decreasing the effects of negative plant– soil feedback (Zuppinger-Dingley et al., 2016). However, in our study the lower performance of selected phytometers in monocultures suggests that the effects of pathogens in the soil are in fact stronger because of the negative plant–soil feedback developed over time (Schnitzer et al., 2011; Thakur et al., 2021). Given the higher performance of *naïve* phytometers in low- diversity communities, new plants transplanted into an “unknown” soil and its associated community were apparently less affected by these dynamics. However, highly diverse communities are characterized by strong competition for light and resources. This may have caused the shift in the observed pattern, leading to a greater performance of *selected* phytometers and supporting the view that plant–soil feedbacks could vary dependent on interspecific competition (Lekberg et al., 2018). As above, the advantage of *selected* phytometers in high-diversity communities could additionally be explained by evolutionary processes. These processes likely led *selected* phytometers to adapt to the plant communities of their original environment and promoted greater performance. These findings add to previous results that selection in high- vs. low-diversity communities leads to genetic modification of plant phenotypes, resulting in plants with higher performance when re-planted in their “home-diversity” environment than wen planted into an “away-diversity” environment (Dietrich et al., 2021; van Moorsel et al., 2018).

Overall, our results show that 17 years of selection history in different biodiversity environments induced measurable plant phenotypic responses to plant and soil community diversity and history. Negative plant–soil feedbacks, typical of low-diversity communities, significantly reduced the performance of phytometers that underwent selection in monocultures. Conversely, adaptive responses could be measured as increased plant performance and trait expression in high-diversity communities, possibly due to adaptation to biotic environments characterized by different levels of plant species richness and associated soil communities.

## Conclusions

After accounting for competition for light, we observed a higher performance of plants in their original environments in highly diverse communities. Therefore, we confirm our first hypothesis (H1) that plants show higher performance in their original environment. We also found that selection in high- vs. low-diversity communities led to adaptive phenotypic responses, largely driven by plant community history effects and plant–soil feedbacks. Therefore, we confirm our second hypothesis (H2) on adaptation of plants that underwent selective processes. We conclude that selection history in communities of different diversity led to adaptive phenotypic responses, promoted by interactions with both plant and soil communities. To better understand co-evolutionary processes between plant and soil communities, we encourage more field studies investigating both the direct roles of plant community history and soil community on selected vs. unselected plants.

## Acknowledgements

We thank the gardeners, technicians and student helpers for maintaining the field site and Anne Ebeling for coordinating the Jena Experiment. We are grateful to Petra Hoffmann for plant chemical analyses, Konstantin Albrecht for help with seed preparation, the technicians of the experimental field station Bad Lauchstädt for help with plant cultivation and the students helpers involved in the projects for their help in the field and in the lab. We thank the German Research Foundation (RO2397-10, DU404-15 in FOR 5000) for funding.

## Author contributions

Francesca De Giorgi, Christiane Roscher and Walter Durka conceived the idea, designed the study and established the phytometer experiment. Francesca De Giorgi performed field and lab work. Francesca De Giorgi and Yuanyuan Huang analyses the data with the support of Christiane Roscher. Francesca De Giorgi drafted the manuscript. All authors contributed to the discussion and revision of the manuscript.

## Data availability statement

This work is based on data elaborated by the subproject 10 “Plant trait variation and evolution in the biodiversity- ecosystem functioning context” of the Jena Experiment. The datatsets will be publicly available in the Jena Experiment database (https://jexis.idiv.de/) with the publication of the manuscript.

## Conflict of interest

The authors declare no conflict of interest.

## Notes

### Competing Interest Statement

The authors have declared no competing interest.

