## Supplementary_material for "Selection shapes plant performance in a grassland biodiversity experiment"

Table S1: Overview of the experimental design of the phytometer experiment showing the species used in the experiment, the number of plots per species-richness level in which they were transplanted and where they belong to the sown composition (in brackets), the planting dates and the number of phytometers for each species.

| Species | Species-richness level | | | | | | Planting date | No. phytometers |
| --- | --- | --- | --- | --- | --- | --- | --- | --- |
|  | 1 | 2 | 4 | 8 | 16 | 60 |  |  |
| *Geranium pratense* | 1(1) |  | 1(1) | 1(1) | 4(7) | 1(4) | 12-15/04/20 | 269 |
| *Ranunculus acris* |  | 2(2) |  | 2(2) | 3(3) | 1(4) | 09/04/20 | 287 |
| *Crepis biennis* | 1(1) |  | 2(2) |  | 2(3) | 1(4) | 02/10/20 | 256 |
| *Plantago lanceolata* | 1(1) | 2(3) | 2(5) | 2(2) | 2(3) | 1(4) | 08-09/04/20 | 567 |
| *Plantago media* |  | 1(1) | 2(2) | 3(3) | 3(4) | 1(4) | 09-10/04/20 | 432 |
| *Lotus corniculatus* |  | 1(1) | 1(1) | 4(4) | 3(4) | 1(4) | 14/04/20 | 453 |
| *Medicago x varia* | 1(1) | 1(1) | 2(2) | 1(1) | 3(3) | 1(4) | 09-14/04/20 | 393 |
| *Alopecurus pratensis* |  | 1(1) | 0(1) | 2(2) | 4(4) | 1(4) | 04-05/04/20 | 384 |
| *Trisetum flavescens* |  | 2(2) | 1(1) | 3(3) | 3(3) | 1(4) | 06-07/04/20 | 480 |

Table S2: Summary of the analysis of Community History Experiment using linear models. Shown are the degrees of freedom, mean square values and calculation of F values for each explanatory variable. Degrees of freedom (DF) refer to n = 1509 individuals, which differ according to the variable and the time of measurements.

| Source of variation | DF | Mean Square | F |
| --- | --- | --- | --- |
| Block (B) | 3 | MS_B_ | MS_B_/MS_P_ |
| Species richness (SR) log-scale | 1 | MS_SR_ | MS_SR_/MS_P_ |
| Treatment (T) | 2 | MS_T_ | MS_T_/MS_SP_ |
| SR x T | 2 | MS_SR x T_ | MS_SR x T_/MS_SP_ |
| Functional group identity (FG-ID) | 3 | MS_FG-ID_ | MS_FG-ID_/MS_P_ |
| SR x FG-ID | 3 | MS_SR x FG-ID_ | MS_SR x FG-ID_/MS_Sp-ID x P_ |
| T x FG-ID | 6 | MS_T x FG-ID_ | MS_T x FG-ID_/ MS_Sp-ID x P_ |
| Species identity (Sp-ID) | 5 | MS_Sp-ID_ | MS_Sp-ID_/ MS_Sp-ID x P_ |
| SR x Sp-ID | 5 | MS_SR x Sp-ID_ | MS_SR x Sp-ID_/ MS_Sp-ID x P_ |
| T x Sp-ID | 10 | MS_T x Sp-ID_ | MS_T x Sp-ID_/MS_SP_ |
| Plot | 47 | MS_P_ | MS_P_/MS_SP_ |
| Sp-ID x Plot | 8 | MS_Sp-ID x P_ | MS_Sp-ID x P_/MS_SP_ |
| Subplot | 100 | MS_SP_ | MS_SP_/MS_Sp-ID x SP_ |
| Sp-ID x Subplot | 28 | MS_Sp-ID x SP_ | MS_Sp-ID x SP_/ MS_T x SF_ |
| Seed family | 210 | MS_SF_ | MS_SF_/ MS_T x SF_ |
| T x Seed family | 307 | MS_T x SF_ | MS_T x SF_/MS_R_ |
| Residuals | 768 | MS_R_ |  |

Table S3: Summary of the analysis of the Selection Experiment using linear models. Shown are the degree of freedom, mean square values and calculation of F value for each explanatory variable. Degrees of freedom (DF) refer to n = 951 individuals, which differ according to the variable and the time of measurements.

| Source of variation | DF | Mean Square | F value |
| --- | --- | --- | --- |
| Block (B) | 3 | MS_B_ | MS_B_/MS_P_ |
| Species richness (SR) log-scale | 1 | MS_SR_ | MS_SR_/MS_P_ |
| Selection (SL) | 1 | MS_SL_ | MS_SL_/MS_SL X Sp-ID_ |
| SR x SL | 1 | MS_SR x SL_ | MS_SR x SL_/MS_SL X P_ |
| Functional group identity (FG-ID) | 3 | MS_FG-ID_ | MS_FG-ID_/MS_P_ |
| SR x FG-ID | 3 | MS_SR x FG-ID_ | MS_SR x FG-ID_/MS_Sp-ID x P_ |
| SL x FG-ID | 3 | MS_SL x FG-ID_ | MS_SL x FG-ID_/ MS_Sp-ID x P_ |
| Species identity (Sp-ID) | 5 | MS_Sp-ID_ | MS_Sp-ID_/ MS_Sp-ID x P_ |
| SR x Sp-ID | 5 | MS_SR x Sp-ID_ | MS_SR x Sp-ID_/ MS_Sp-ID x P_ |
| SL x Sp-ID | 5 | MS_SL x Sp-ID_ | MS_SL x Sp-ID_/MS_SF_ |
| Plot (P) | 47 | MS_P_ | MS_P_/MS_Sp-ID x P_ |
| Sp-ID x Plot | 7 | MS_Sp-ID x P_ | MS_Sp-ID x P_/MS_SF_ |
| SL x Plot | 48 | MS_SL X P_ | MS_SP_/MS_SF_ |
| Seed family | 204 | MS_SF_ | MS_SF_/ MS_R_ |
| Residuals | 614 | MS_R_ |  |

Table S4: Results of linear models of the *Community History Experiment* testing effects of canopy height of the surrounding vegetation, sown species richness (SR), treatment, their interaction, identity of the functional group (FG-ID), its interaction with species richness and treatment, species and its interaction with species richness and treatment on plant performance and trait expression. If variables were measured at different time points, it is indicated with t1= summer 2020, t2= spring 2021, and t3= summer 2021. Shown are F and P values. Abbreviations of variable names: LeafG= leaf greenness, C_Leaf_ = leaf carbon concentration, N_Leaf_ = leaf nitrogen concentration, SLA = specific leaf area.

| Source of variation | Plant biomass | | RGR_t3-t1 | | | | | | Plant height_t1 | | | | | | Plant height_t2 | | | | | | Plant height_t3 | | |
| --- | --- | --- | --- | --- | --- | --- | --- | --- | --- | --- | --- | --- | --- | --- | --- | --- | --- | --- | --- | --- | --- | --- | --- |
|  | F | P | | F | | P | | | | F | | P | | | | F | | P | | | | F | P |
| Canopy height | 5.33 | 0.070 | | 189.75 | | 0.001 | | | | 19.23 | | 0.002 | | | | 31.79 | | <0.001 | | | | 26.36 | <0.001 |
| Block | 0.27 | 0.603 | | 1.83 | | 0.185 | | | | 59.38 | | <0.001 | | | | 284.94 | | <0.001 | | | | 47.86 | <0.001 |
| Species richness (SR) (log) | 35.18 | <0.001 | | 16.37 | | 0.001 | | | | 13.25 | | 0.001 | | | | 19.88 | | <0.001 | | | | 9.33 | 0.006 |
| Treatment | 3.42 | 0.040 | | 0.01 | | 0.991 | | | | 0.32 | | 0.726 | | | | 4.38 | | 0.015 | | | | 1.63 | 0.207 |
| SR x Treatment | 1.09 | 0.342 | | 1.74 | | 0.190 | | | | 1.13 | | 0.328 | | | | 0.96 | | 0.387 | | | | 1.77 | 0.180 |
| FG-ID | 27.79 | <0.001 | | 6.53 | | 0.009 | | | | 17.84 | | <0.001 | | | | 53.87 | | <0.001 | | | | 56.70 | <0.001 |
| SR x FG-ID | 1.50 | 0.500 | | 891.00 | | 0.021 | | | | 13.20 | | 0.005 | | | | 3.05 | | 0.092 | | | | 4.33 | 0.322 |
| Treatment x FG-ID | 3.50 | 0.379 | | 4.00 | | 0.349 | | | | 1.18 | | 0.425 | | | | 1.07 | | 0.453 | | | | 0.92 | 0.645 |
| Species-ID | 34.33 | 0.125 | | 2280.00 | | 0.015 | | | | 25.65 | | 0.001 | | | | 40.27 | | <0.001 | | | | 5.44 | 0.303 |
| SR x Species-ID | 0.50 | 0.707 | | 66.00 | | 0.087 | | | | 7.20 | | 0.018 | | | | 3.93 | | 0.043 | | | | 0.50 | 0.707 |
| Treatment x Species-ID | 1.59 | 0.180 | | 0.40 | | 0.810 | | | | 0.86 | | 0.549 | | | | 1.61 | | 0.113 | | | | 1.65 | 0.162 |
| Plot | 2.20 | 0.009 | | 1.74 | | 0.087 | | | | 2.50 | | <0.001 | | | | 2.90 | | <0.001 | | | | 3.13 | <0.001 |
| Species-ID x Plot | 0.27 | 0.603 | | 0.01 | | 0.925 | | | | 1.07 | | 0.384 | | | | 1.20 | | 0.309 | | | | 0.89 | 0.351 |
| Subplot | 2.92 | 0.055 | | 22.20 | | <0.001 | | | | 1.53 | | 0.137 | | | | 1.16 | | 0.337 | | | | 1.43 | 0.312 |
| Species-ID x Subplot | 1.25 | 0.273 | | 0.12 | | 0.994 | | | | 1.98 | | 0.008 | | | | 1.90 | | 0.005 | | | | 1.78 | 0.084 |
| SF | 1.61 | 0.003 | | 1.10 | | 0.324 | | | | 1.13 | | 0.166 | | | | 1.12 | | 0.173 | | | | 1.17 | 0.173 |
| Treatment x SF | 0.76 | 0.984 | | 0.98 | | 0.551 | | | | 1.02 | | 0.403 | | | | 1.43 | | <0.001 | | | | 0.90 | 0.791 |
| Source of variation | LeafG_t1 | | LeafG_t3 | | | | SLA | | | | | | N_Leaf_ | | | | | | C_Leaf_ | | | | |
|  | F | P | | F | P | | | F | | | P | | | F | | | P | | | F | | | P |
| Canopy height | 9.80 | 0.010 | | 42.61 | 0.002 | | | 34,1 | | | <0.001 | | | 30.57 | | | <0.001 | | | 7.73 | | | <0.001 |
| Block | 66.37 | <0.001 | | 23.84 | <0.001 | | | 303,5 | | | <0,001 | | | 0.00 | | | 1.000 | | | 8.85 | | | 0.004 |
| Species richness (SR) (log) | 38.50 | <0.001 | | 44.15 | <0.001 | | | 9,7 | | | 0,003 | | | 17.59 | | | <0.001 | | | 2.42 | | | 0.126 |
| Treatment | 0.84 | 0.434 | | 1.45 | 0.243 | | | 4,5 | | | 0,013 | | | 0.00 | | | 1.000 | | | 2.08 | | | 0.130 |
| SR x Treatment | 0.16 | 0.849 | | 2.08 | 0.136 | | | 1,2 | | | 0,313 | | | 2.40 | | | 0.095 | | | 0.26 | | | 0.771 |
| FG-ID | 101.88 | <0.001 | | 90.99 | <0.001 | | | 70,8 | | | <0,001 | | | 206.89 | | | <0.001 | | | 67.69 | | | <0.001 |
| SR x FG-ID | 8.81 | 0.013 | | 0.53 | 0.698 | | | 1,0 | | | 0,439 | | | 2.22 | | | 0.163 | | | 2.05 | | | 0.185 |
| Treatment x FG-ID | 1.12 | 0.446 | | 2.50 | 0.439 | | | 0,3 | | | 0,931 | | | 0.11 | | | 0.992 | | | 0.72 | | | 0.647 |
| Species-ID | 6.60 | 0.022 | | 26.38 | 0.142 | | | 15,6 | | | 0,001 | | | 28.27 | | | <0.001 | | | 13.54 | | | 0.001 |
| SR x Species-ID | 4.71 | 0.046 | | 2.65 | 0.398 | | | 1,1 | | | 0,440 | | | 1.33 | | | 0.341 | | | 1.11 | | | 0.426 |
| Treatment x Species-ID | 3.16 | 0.003 | | 0.70 | 0.627 | | | 1,8 | | | 0,066 | | | 0.72 | | | 0.703 | | | 1.04 | | | 0.415 |
| Plot | 3.09 | <0.001 | | 2.43 | 0.004 | | | 2,1 | | | 0,001 | | | 3.83 | | | <0.001 | | | 2.58 | | | <0.001 |
| Species-ID x Plot | 1.23 | 0.300 | | 0.21 | 0.652 | | | 3,4 | | | 0,002 | | | 3.61 | | | 0.001 | | | 1.69 | | | 0.109 |
| Subplot | 3.04 | 0.004 | | 1.59 | 0.249 | | | 1,6 | | | 0,072 | | | 1.02 | | | 0.497 | | | 0.58 | | | 0.971 |
| Species-ID x Subplot | 0.59 | 0.913 | | 1.15 | 0.332 | | | 1,0 | | | 0,427 | | | 1.72 | | | 0.014 | | | 1.76 | | | 0.011 |
| SF | 0.93 | 0.726 | | 1.11 | 0.263 | | | 1,1 | | | 0,329 | | | / | | | / | | | / | | | / |
| Treatment x SF | 1.08 | 0.187 | | 1.23 | 0.049 | | | 0,9 | | | 0,843 | | | / | | | / | | | / | | | / |

Table S5: Results of linear models and generalized mixed effect models of the *Community History Experiment* testing effects of sown species richness (SR), treatment, their interaction, identity of the functional group (FG-ID), its interaction with species richness and treatment, species and its interaction with species richness and treatment on trait performance and trait expression. If variables were measured at different time points, it is indicated with t1= summer 2020 and t2= spring 2021. Shown are F and P values. Abbreviations of variable names: LeafG= leaf greenness.

| Source of variation | Survival_t1 | | Survival_t2 | | Plant height_t1 | | Plant height_t2 | | LeafG_t1 | |
| --- | --- | --- | --- | --- | --- | --- | --- | --- | --- | --- |
|  | χ2 | P | χ2 | P | F | P | F | P | F | P |
| Block | 7.62 | 0.055 | 13.48 | 0.004 | 7.29 | <0.001 | 13.30 | <0.001 | 3.46 | 0.024 |
| Species richness (SR) (log) | 3.40 | 0.065 | 0.37 | 0.545 | 5.60 | 0.022 | 3.20 | 0.080 | 15.51 | <0.001 |
| Treatment | 3.18 | 0.204 | 12.57 | 0.002 | 0.99 | 0.376 | 14.42 | <0.001 | 1.56 | 0.215 |
| SR x Treatment | 0.72 | 0.698 | 3.63 | 0.163 | 1.71 | 0.187 | 1.39 | 0.254 | 0.41 | 0.666 |
| FG-ID | 12.47 | 0.006 | 6.76 | 0.080 | 29.48 | <0.001 | 75.72 | <0.001 | 90.40 | <0.001 |
| SR x FG-ID | 10.13 | 0.018 | 14.28 | 0.003 | 9.35 | 0.011 | 2.47 | 0.137 | 11.38 | 0.007 |
| Treatment x FG-ID | 10.88 | 0.092 | 5.00 | 0.544 | 0.83 | 0.589 | 1.00 | 0.485 | 2.14 | 0.189 |
| Species-ID | 9.73 | 0.045 | 13.31 | 0.021 | 25.89 | 0.001 | 40.48 | <0.001 | 12.21 | 0.005 |
| SR x Species-ID | 12.24 | 0.016 | 3.80 | 0.578 | 7.73 | 0.015 | 3.32 | 0.064 | 6.22 | 0.025 |
| Treatment x Species-ID | 11.42 | 0.179 | 19.90 | 0.030 | 0.85 | 0.558 | 2.14 | 0.028 | 2.87 | 0.007 |
| Plot | / | / | / | / | 2.92 | <0.001 | 3.29 | <0.001 | 3.65 | <0.001 |
| Species-ID x Plot | / | / | / | / | 1.38 | 0.231 | 1.39 | 0.211 | 0.81 | 0.564 |
| Subplot | / | / | / | / | 0.91 | 0.643 | 1.03 | 0.484 | 1.96 | 0.028 |
| Species-ID x Subplot | / | / | / | / | 3.00 | <0.001 | 2.11 | 0.001 | 0.95 | 0.540 |
| SF | / | / | / | / | 1.15 | 0.130 | 1.11 | 0.187 | 0.90 | 0.802 |
| Treatment x SF | / | / | / | / | 1.03 | 0.366 | 1.44 | <0.001 | 1.07 | 0.199 |

Table S6: Results of linear models for the *Selection Experiment* testing effects of canopy height of the surrounding vegetation height, sown species richness (SR), selection history, their interaction, identity of the functional group (FG-ID), its interaction with species richness and selection history, species and its interaction with species richness and selection history on trait performance and trait expression. If variables were measured at different time points, it is indicated with t1= summer 2020, t2= spring 2021, and t3= summer 2021. Shown are F and P values. Abbreviations of variable names: LeafG= leaf greenness, C_Leaf_ = leaf carbon concentration, N_Leaf_ = leaf nitrogen concentration, SLA = specific leaf area.

| Source of variation | Plant biomass | | RGR_t3-t1 | | | | Plant height_t1 | | | | | Plant height_t2 | | | | | Plant height_t3 | | | | |
| --- | --- | --- | --- | --- | --- | --- | --- | --- | --- | --- | --- | --- | --- | --- | --- | --- | --- | --- | --- | --- | --- |
|  | F | P | | F | P | | | F | | P | | | F | | P | | | F | | P | |
| Canopy height | 1.54 | 0.231 | | 0.65 | 0.596 | | | 7.22 | | <0.001 | | | 7.87 | | <0.001 | | | 33.72 | | <0.001 | |
| Block | 0.52 | 0.477 | | 1.23 | 0.282 | | | 104.16 | | <0.001 | | | 207.14 | | <0.001 | | | 28.81 | | <0.001 | |
| Species richness (SR) (log) | 12.30 | 0.002 | | 2.69 | 0.123 | | | 6.79 | | 0.012 | | | 13.34 | | 0.001 | | | 3.58 | | 0.072 | |
| Selection | 2.67 | 0.201 | | 0.86 | 0.452 | | | 0.57 | | 0.492 | | | 0.00 | | 1.000 | | | 6.94 | | 0.078 | |
| SR x Selection | 0.52 | 0.477 | | 3.70 | 0.071 | | | 1.40 | | 0.242 | | | 0.31 | | 0.580 | | | 1.43 | | 0.243 | |
| FG-ID | 16.26 | <0.001 | | 1.63 | 0.232 | | | 6.77 | | 0.001 | | | 33.27 | | <0.001 | | | 46.16 | | <0.001 | |
| SR x FG-ID | 13.50 | 0.189 | | 49.00 | 0.090 | | | 6.78 | | 0.024 | | | 1.14 | | 0.399 | | | 9.50 | | 0.224 | |
| Selection x FG-ID | 0.50 | 0.707 | | 7.25 | 0.254 | | | 1.15 | | 0.402 | | | 0.57 | | 0.654 | | | 26.50 | | 0.136 | |
| Species-ID | 17.33 | 0.174 | | 40.50 | 0.110 | | | 10.32 | | 0.007 | | | 24.63 | | <0.001 | | | 133.00 | | 0.064 | |
| SR x Species-ID | 0.50 | 0.707 | | 2.25 | 0.426 | | | 3.97 | | 0.066 | | | 3.14 | | 0.084 | | | 1.50 | | 0.500 | |
| Species-ID x Selection | 1.99 | 0.120 | | 1.44 | 0.244 | | | 2.00 | | 0.095 | | | 1.29 | | 0.271 | | | 1.33 | | 0.267 | |
| Plot | 6.91 | 0.293 | | 11.54 | 0.227 | | | 1.96 | | 0.202 | | | 1.79 | | 0.215 | | | 32.00 | | 0.139 | |
| Species-ID x Plot | 0.66 | 0.417 | | 0.41 | 0.523 | | | 3.75 | | 0.001 | | | 3.09 | | 0.004 | | | 0.17 | | 0.684 | |
| Plot x Selection | 1.27 | 0.198 | | 0.83 | 0.650 | | | 1.36 | | 0.069 | | | 1.88 | | 0.001 | | | 1.75 | | 0.026 | |
| SF | 1.22 | 0.095 | | 1.18 | 0.180 | | | 1.12 | | 0.130 | | | 1.38 | | 0.001 | | | 0.80 | | 0.915 | |
| Source of variation | LeafG_t1 |  | | Leaf G_t3 | |  | | | SLA | | | | | N_Leaf_ | | | | | C_Leaf_ | | |
|  | F | P | | F | | P | | | F | | P | | | F | | P | | | F | | P |
| Canopy height | 1.14 | 0.345 | | 5.38 | | 0.006 | | | 11.68 | | <0.001 | | | 16.36 | | <0.001 | | | 4.00 | | 0.013 |
| Block | 78.72 | <0.001 | | 2.30 | | 0.142 | | | 268.55 | | <0.001 | | | 3.27 | | 0.077 | | | 1.59 | | 0.213 |
| Species richness (SR) (log) | 35.92 | <0.001 | | 33.87 | | <0.001 | | | 5.20 | | 0.027 | | | 31.79 | | <0.001 | | | 10.80 | | 0.002 |
| Selection | 0.49 | 0.522 | | 0.88 | | 0.418 | | | 0.18 | | 0.687 | | | 0.83 | | 0.403 | | | 2.92 | | 0.148 |
| SR x Selection | 0.43 | 0.517 | | 0.51 | | 0.484 | | | 2.34 | | 0.133 | | | 4.90 | | 0.032 | | | 3.43 | | 0.070 |
| FG-ID | 58.72 | <0.001 | | 69.74 | | <0.001 | | | 44.09 | | <0.001 | | | 138.00 | | <0.001 | | | 38.07 | | <0.001 |
| SR x FG-ID | 6.83 | 0.023 | | 20.33 | | 0.155 | | | 1.02 | | 0.439 | | | 7.78 | | 0.012 | | | 0.37 | | 0.778 |
| Selection x FG-ID | 1.55 | 0.295 | | 12.00 | | 0.200 | | | 0.72 | | 0.569 | | | 1.56 | | 0.283 | | | 3.19 | | 0.093 |
| Species-ID | 7.89 | 0.014 | | 36.93 | | 0.120 | | | 11.03 | | 0.003 | | | 69.07 | | <0.001 | | | 14.59 | | 0.001 |
| SR x Species-ID | 3.26 | 0.095 | | 17.22 | | 0.168 | | | 0.92 | | 0.519 | | | 3.27 | | 0.077 | | | 9.28 | | 0.005 |
| Species-ID x Selection | 3.96 | 0.004 | | 0.97 | | 0.411 | | | 1.69 | | 0.138 | | | 3.14 | | 0.009 | | | 0.42 | | 0.835 |
| Plot | 2.01 | 0.194 | | 13.74 | | 0.210 | | | 0.90 | | 0.629 | | | 3.38 | | 0.048 | | | 3.68 | | 0.038 |
| Species-ID x Plot | 2.38 | 0.030 | | 0.12 | | 0.724 | | | 4.43 | | <0.001 | | | 1.12 | | 0.348 | | | 0.47 | | 0.853 |
| Plot x Selection | 1.13 | 0.276 | | 0.91 | | 0.596 | | | 1.15 | | 0.243 | | | 1.60 | | 0.009 | | | 1.43 | | 0.037 |
| SF | 1.11 | 0.148 | | 1.85 | | <0.001 | | | 0.94 | | 0.717 | | | / | | / | | | / | | / |

Table S7: Results of linear models and generalized mixed effect models of the *Selection Experiment* testing effects of sown species richness (SR), treatment, their interaction, identity of the functional group (FG-ID), its interaction with species richness and treatment, species and its interaction with species richness and treatment on trait performance and trait expression. If variables were measured at different time points, it is indicated with t1= summer 2020 and t2= spring 2021. Shown are F and P values. Abbreviations of variable names: LeafG= leaf greenness.

| Source of variation | Survival_t1 | | Survival_t2 | | | Plant height_t1 | | | Plant height_t2 | | | LeafG_t1 | | |
| --- | --- | --- | --- | --- | --- | --- | --- | --- | --- | --- | --- | --- | --- | --- |
|  | χ2 | P | | χ2 | P | | F | P | | F | P | | F | P |
| Block | 3.62 | 0.305 | | 9.67 | 0.022 | | 5.47 | 0.003 | | 8.54 | <0.001 | | 1.16 | 0.334 |
| Species richness (SR) (log) | 2.17 | 0.140 | | 1.42 | 0.234 | | 2.45 | 0.125 | | 5.16 | 0.028 | | 19.79 | <0.001 |
| Selection | 2.12 | 0.145 | | 0.37 | 0.545 | | 0.67 | 0.460 | | 1.11 | 0.340 | | 1.73 | 0.258 |
| SR x Selection | 0.01 | 0.941 | | 0.98 | 0.321 | | 2.77 | 0.103 | | 0.33 | 0.566 | | 1.05 | 0.311 |
| FG-ID | 6.14 | 0.105 | | 2.19 | 0.535 | | 15.47 | <0.001 | | 47.43 | <0.001 | | 65.84 | <0.001 |
| SR x FG-ID | 4.66 | 0.199 | | 10.00 | 0.019 | | 5.48 | 0.037 | | 0.64 | 0.610 | | 5.76 | 0.034 |
| Selection x FG-ID | 4.03 | 0.258 | | 2.15 | 0.542 | | 1.13 | 0.409 | | 1.23 | 0.362 | | 0.94 | 0.477 |
| Species-ID | 4.69 | 0.321 | | 8.05 | 0.154 | | 11.35 | 0.006 | | 26.98 | <0.001 | | 7.88 | 0.014 |
| SR x Species-ID | 4.45 | 0.349 | | 5.61 | 0.346 | | 4.79 | 0.045 | | 3.20 | 0.070 | | 3.68 | 0.076 |
| Species-ID x Selection | 8.82 | 0.066 | | 13.33 | 0.020 | | 2.36 | 0.054 | | 1.05 | 0.388 | | 1.83 | 0.123 |
| Plot | / | / | | / | / | | 1.90 | 0.215 | | 1.74 | 0.207 | | 2.12 | 0.174 |
| Species-ID x Plot | / | / | | / | / | | 4.07 | 0.001 | | 3.65 | <0.001 | | 2.33 | 0.034 |
| Plot x Selection | / | / | | / | / | | 1.42 | 0.045 | | 1.75 | 0.003 | | 1.40 | 0.052 |
| SF | / | / | | / | / | | 1.09 | 0.177 | | 1.38 | 0.001 | | 1.13 | 0.107 |

Table S8: Results of Tukey’s HSD test used to identify differences among four functional groups in the *Community History Experiment*. Shown are mean values and standard error (SE) of measured values for each functional group; letters indicate statistically significant differences.

| Trait | Grasses | | | Legumes | | | | Small herbs | | | | Tall herbs | | | |
| --- | --- | --- | --- | --- | --- | --- | --- | --- | --- | --- | --- | --- | --- | --- | --- |
|  | Mean | SE | Group | | Mean | SE | Group | | Mean | SE | Group | | Mean | SE | Group |
| Survival rate_t1 | 0.949 | 0.009 | b | | 0.905 | 0.012 | ab | | 0.926 | 0.010 | b | | 0.861 | 0.018 | a |
| Survival rate_t2 | 0.728 | 0.017 | a | | 0.605 | 0.020 | a | | 0.756 | 0.016 | a | | 0.689 | 0.019 | a |
| Survival rate_t3 | 0.616 | 0.019 | ab | | 0.515 | 0.020 | ab | | 0.691 | 0.017 | b | | 0.491 | 0.021 | a |
| Flowering proportion | 0.373 | 0.022 | b | | 0.457 | 0.025 | b | | 0.575 | 0.020 | b | | 0.262 | 0.022 | a |
| Plant biomass (g) | 0.418 | 0.039 | a | | 2.617 | 0.264 | b | | 1.516 | 0.088 | b | | 0.563 | 0.064 | a |
| Relative growth rate t3_t1 | -0.002 | 0.000 | a | | 0.000 | 0.000 | b | | 0.000 | 0.000 | b | | 0.000 | 0.000 | b |
| Plant height_t1 (cm) | 15.657 | 0.300 | bc | | 20.729 | 0.464 | c | | 10.001 | 0.180 | a | | 13.516 | 0.407 | ab |
| Plant height_t2 (cm) | 35.386 | 0.620 | b | | 16.655 | 0.447 | a | | 15.318 | 0.330 | a | | 15.874 | 0.474 | a |
| Plant height_t3 (cm) | 24.535 | 0.600 | b | | 32.080 | 0.936 | b | | 11.994 | 0.289 | a | | 16.245 | 0.547 | a |
| Leaf greennes_t1 | 28.424 | 0.361 | a | | 49.578 | 0.400 | d | | 36.229 | 0.346 | c | | 32.512 | 0.543 | b |
| Leaf greenness_t3 | 24.154 | 0.398 | a | | 47.035 | 0.477 | d | | 33.032 | 0.347 | c | | 28.016 | 0.409 | b |
| SLA (mm^2^_leaf_ g^-1^_leaf_) | 23.527 | 0.270 | b | | 26.304 | 0.469 | c | | 17.465 | 0.297 | a | | 23.446 | 0.311 | b |
| N_Leaf_ | 18.224 | 0.389 | ab | | 43.723 | 0.581 | c | | 16.611 | 0.258 | a | | 20.991 | 0.453 | b |
| C_Leaf_ | 44.784 | 0.122 | b | | 48.137 | 0.281 | c | | 43.147 | 0.076 | a | | 45.450 | 0.144 | b |

Table S9: Results of Tukey’s HSD test used to identify differences among four functional groups in the *Selection Experiment*. Shown are mean values and standard error (SE) of measured values for each functional group; letters indicate statistically significant differences.

| Trait | Grasses | | | Legumes | | | | Small herbs | | | | Tall herbs | | | |
| --- | --- | --- | --- | --- | --- | --- | --- | --- | --- | --- | --- | --- | --- | --- | --- |
|  | Mean | SE | Group | | Mean | SE | Group | | Mean | SE | Group | | Mean | SE | Group |
| Survival rate_t1 | 0.900 | 0.015 | a | | 0.892 | 0.015 | a | | 0.934 | 0.011 | a | | 0.877 | 0.019 | a |
| Survival rate_t2 | 0.653 | 0.023 | a | | 0.589 | 0.024 | a | | 0.697 | 0.021 | a | | 0.662 | 0.022 | a |
| Survival rate_t3 | 0.514 | 0.024 | ab | | 0.464 | 0.024 | ab | | 0.677 | 0.021 | b | | 0.412 | 0.023 | a |
| Flowering proportion | 0.374 | 0.028 | ab | | 0.447 | 0.030 | ab | | 0.583 | 0.025 | b | | 0.247 | 0.024 | a |
| Plant biomass (g) | 0.316 | 0.036 | a | | 1.568 | 0.172 | b | | 1.299 | 0.091 | b | | 0.555 | 0.070 | a |
| Relative growth rate t3_t1 | -0.002 | 0.000 | a | | 0.000 | 0.000 | b | | 0.000 | 0.000 | b | | 0.000 | 0.000 | b |
| Plant height_t1 (cm) | 14.570 | 0.368 | bc | | 20.488 | 0.597 | c | | 9.876 | 0.206 | a | | 13.749 | 0.439 | ab |
| Plant height_t2 (cm) | 33.901 | 0.771 | b | | 16.741 | 0.592 | a | | 16.005 | 0.441 | a | | 15.469 | 0.497 | a |
| Plant height_t3 (cm) | 24.268 | 0.797 | b | | 32.692 | 1.078 | b | | 12.325 | 0.420 | a | | 17.402 | 0.631 | a |
| Leaf greennes_t1 (cm) | 27.079 | 0.461 | a | | 47.947 | 0.543 | c | | 35.817 | 0.418 | b | | 33.244 | 0.663 | b |
| Leaf greenness_t3 | 23.402 | 0.494 | a | | 45.787 | 0.620 | d | | 31.645 | 0.380 | c | | 27.144 | 0.437 | b |
| SLA (mm^2^_leaf_ g^-1^_leaf_) | 24.015 | 0.361 | bc | | 24.971 | 0.434 | c | | 16.957 | 0.293 | a | | 22.990 | 0.372 | b |
| N_Leaf_ | 18.110 | 0.501 | ab | | 42.489 | 0.849 | c | | 16.413 | 0.355 | a | | 20.785 | 0.509 | b |
| C_Leaf_ | 44.863 | 0.150 | b | | 47.453 | 0.386 | c | | 42.924 | 0.092 | a | | 45.556 | 0.209 | b |

Table S10: Results of Tukey’s HSD test used to identify differences in trait value among the three treatments of the ΔBEF Experiment. Shown are mean values and standard error (SE) of measured values for each functional group; letters indicate statistically significant differences.

| Trait | -PH -SH | | | -PH +SH | | | +PH +SH | | |
| --- | --- | --- | --- | --- | --- | --- | --- | --- | --- |
|  | Mean | SE | Group | Mean | SE | Group | Mean | SE | Group |
| Survival_t2 | 0.687 | 0.016 | a | 0.750 | 0.015 | b | 0.660 | 0.016 | a |
| Survival_t3 | 0.589 | 0.017 | ab | 0.638 | 0.017 | b | 0.533 | 0.017 | a |
| Plant height_t2 (cm) | 22.948 | 0.560 | b | 19.366 | 0.504 | a | 20.673 | 0.547 | a |
| SLA (mm^2^_leaf_ g^-1^_leaf_) | 23.160 | 0.363 | b | 21.664 | 0.309 | a | 21.684 | 0.279 | a |
| N_Leaf_ | 24.697 | 0.767 | b | 24.362 | 0.753 | ab | 23.437 | 0.736 | a |
| C_Leaf_ | 45.453 | 0.193 | a | 45.038 | 0.179 | a | 45.201 | 0.163 | a |

| Trait | -PH -SH | | | -PH +SH | | | +PH +SH | | |
| --- | --- | --- | --- | --- | --- | --- | --- | --- | --- |
|  | Mean | SE | Group | Mean | SE | Group | Mean | SE | Group |
| Canopy height_t1 | 19.024 | 0.389 | a | 17.809 | 0.389 | a | 17.992 | 0.398 | a |
| Canopy height_t2 | 37.985 | 0.673 | a | 32.396 | 0.627 | b | 33.182 | 0.600 | b |
| Canopy height_t3 | 26.130 | 0.441 | a | 24.098 | 0.488 | a | 25.737 | 0.567 | a |

Table S11: Results of Tukey’s HSD test used to identify differences in canopy height among treatments of the ΔBEF Experiment. Shown are mean values and standard error (SE) for canopy height measured at different time points, i.e. t1 = summer 2020, t2 = spring 2021, t3 = summer 2021. Letters indicate statistically significant differences.

Figure S1: Effects of sown species richness on (A) plant height (summer 2020) and (B) leaf greenness (summer 2020) for the *Community History Experiment*. Solid colored lines represent a significant relationship at the functional-group level, while the solid black line represents the mean response across all species.

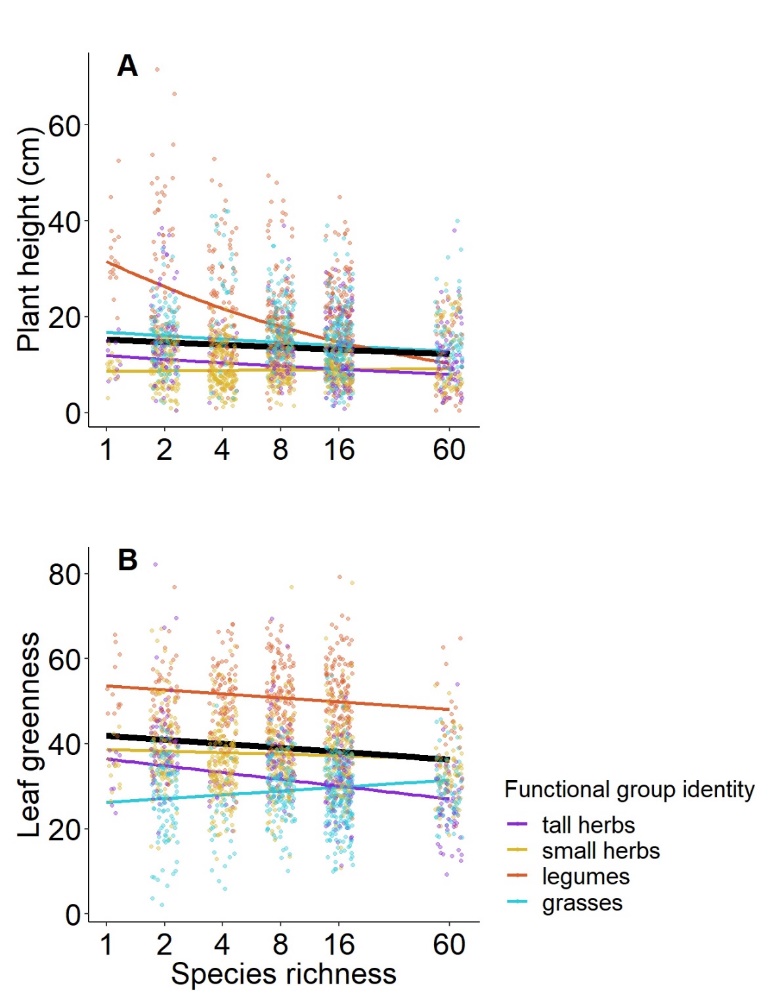

Figure S2: Effects of sown species richness on (A) plant individual biomass (summer 2021), (B) flowering rate (year 2021), (C) leaf greenness (summer 2021), (D) specific leaf area (SLA) (spring 2021), (E) leaf nitrogen concentration (N_Leaf_) (spring 2021), and (F) leaf carbon concentration (C_Leaf_) (spring 2021) for the *Selection Experiment*. Solid colored lines represent a significant relationship at the functional-group level, while the solid black line represents the mean response across all species.

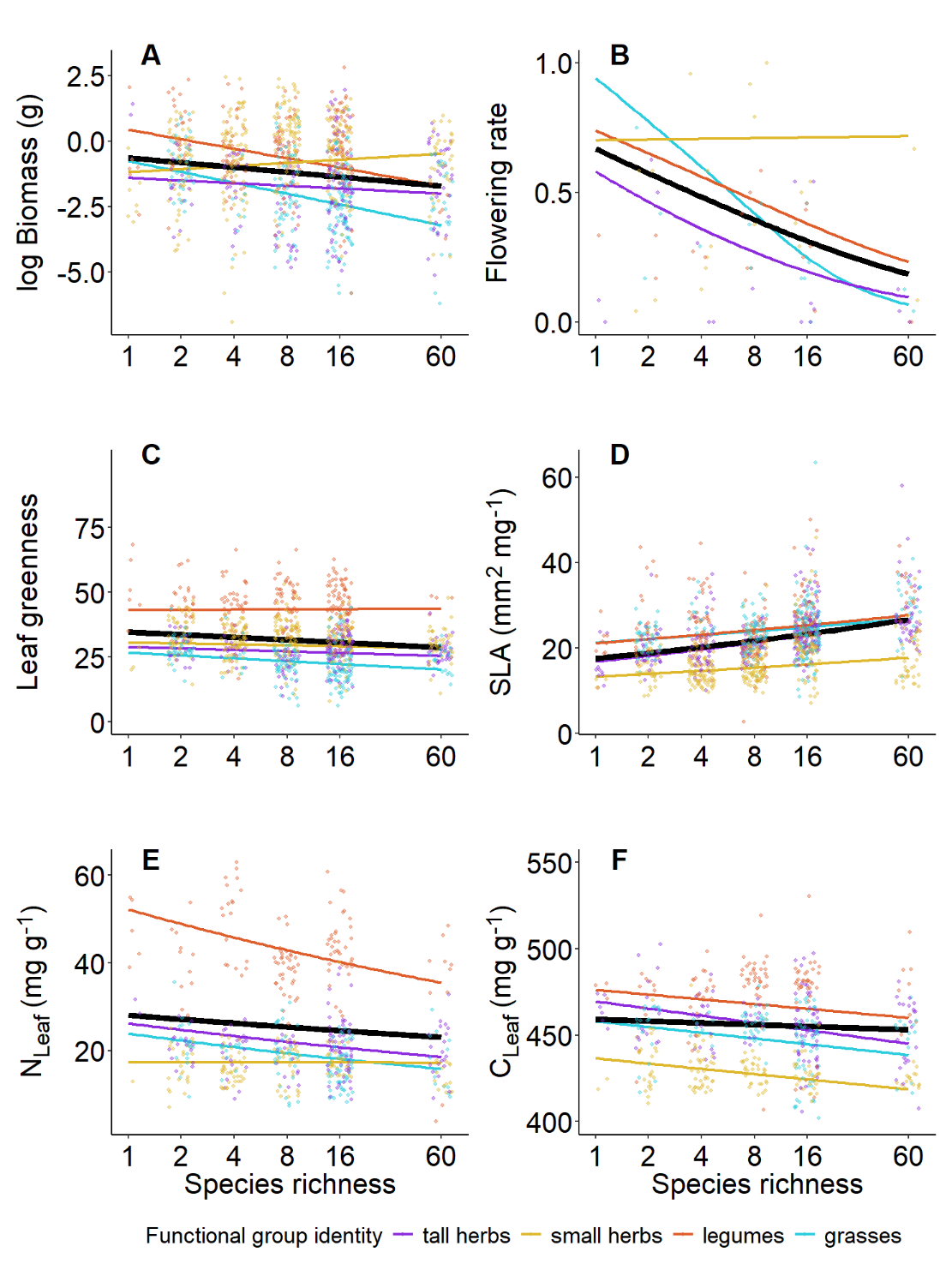

Figure S3: Effects of sown species richness on (A) leaf greenness (summer 2020) and (B) plant height (spring 2021) for the *Selection Experiment*. Solid colored lines represent a significant relationship at the functional-group level, while the solid black line represents the mean response across all species.

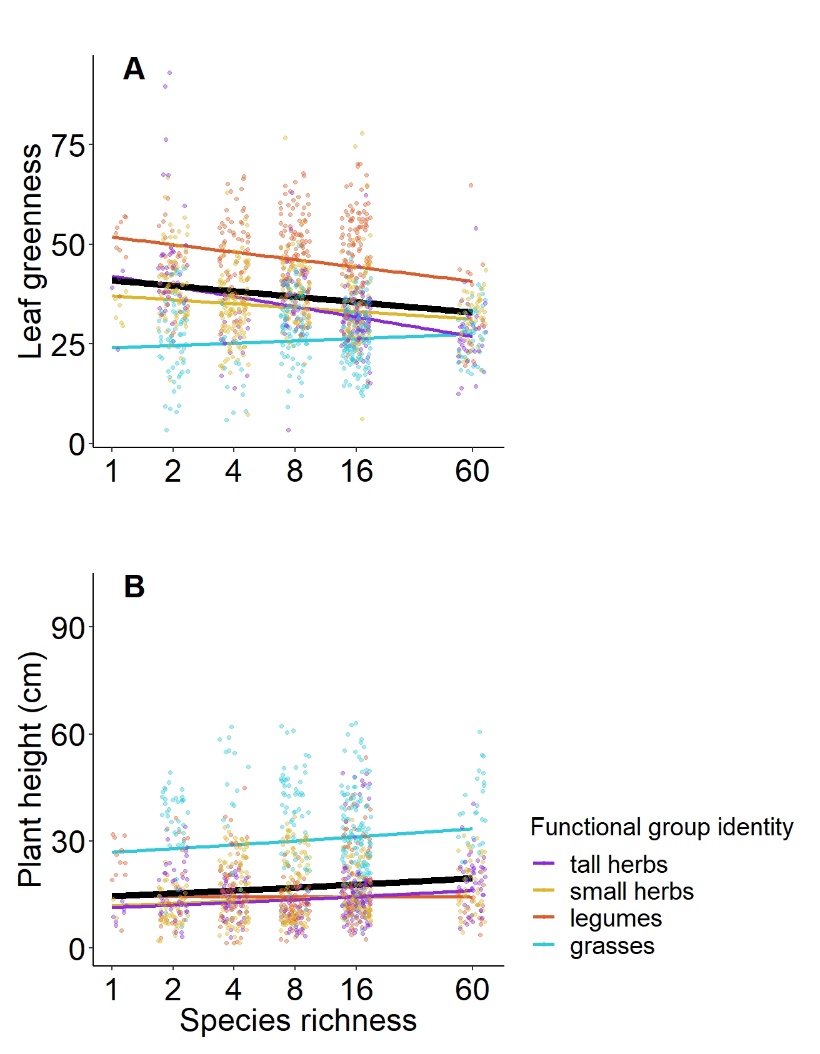

Figure S4: Effects of the treatment environments on (A) survival rate (spring 2021), (B) plant height (summer 2021) and (C) plant aboveground biomass (summer 2021) for the *Community History Experiment*. Solid colored lines represent a significant relationship at the treatment level. Shown are the different treatment environments of the ΔBEF Experiment. i.e. “without plant and soil history” (-PH-SH). “without plant history, with soil history” (-PH+SH) and “with plant and soil history” (+PH+SH).

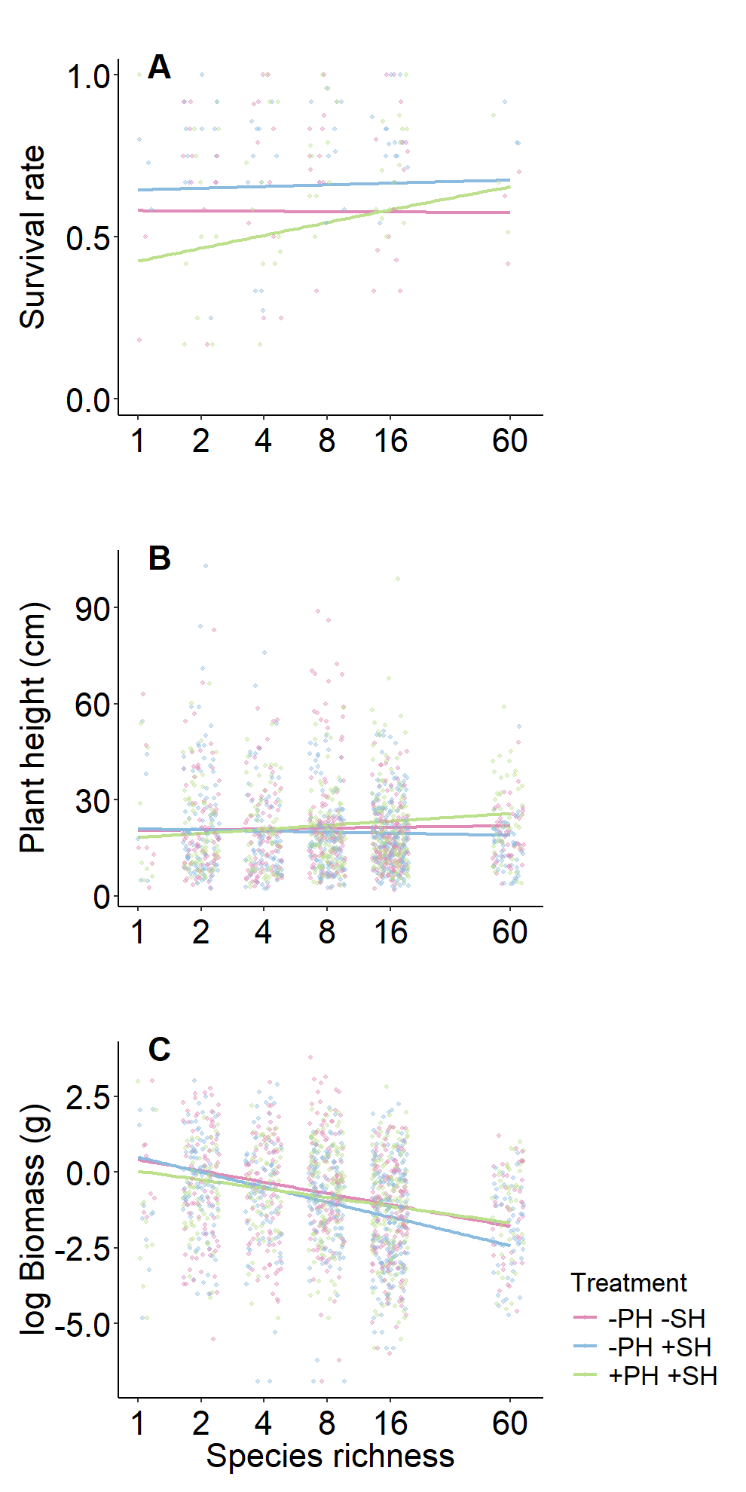

Figure S5: Plot-means of canopy height for the three ΔBEF treatments measured (A) in summer 2020, (C) spring 2021, and (E) summer 2021, and subplot-means of vegetation height between the areas in which phytometers with and without section history were transplanted (right panels) measured (B) in summer 2020, (D) spring 2021, and (F) summer 2021.

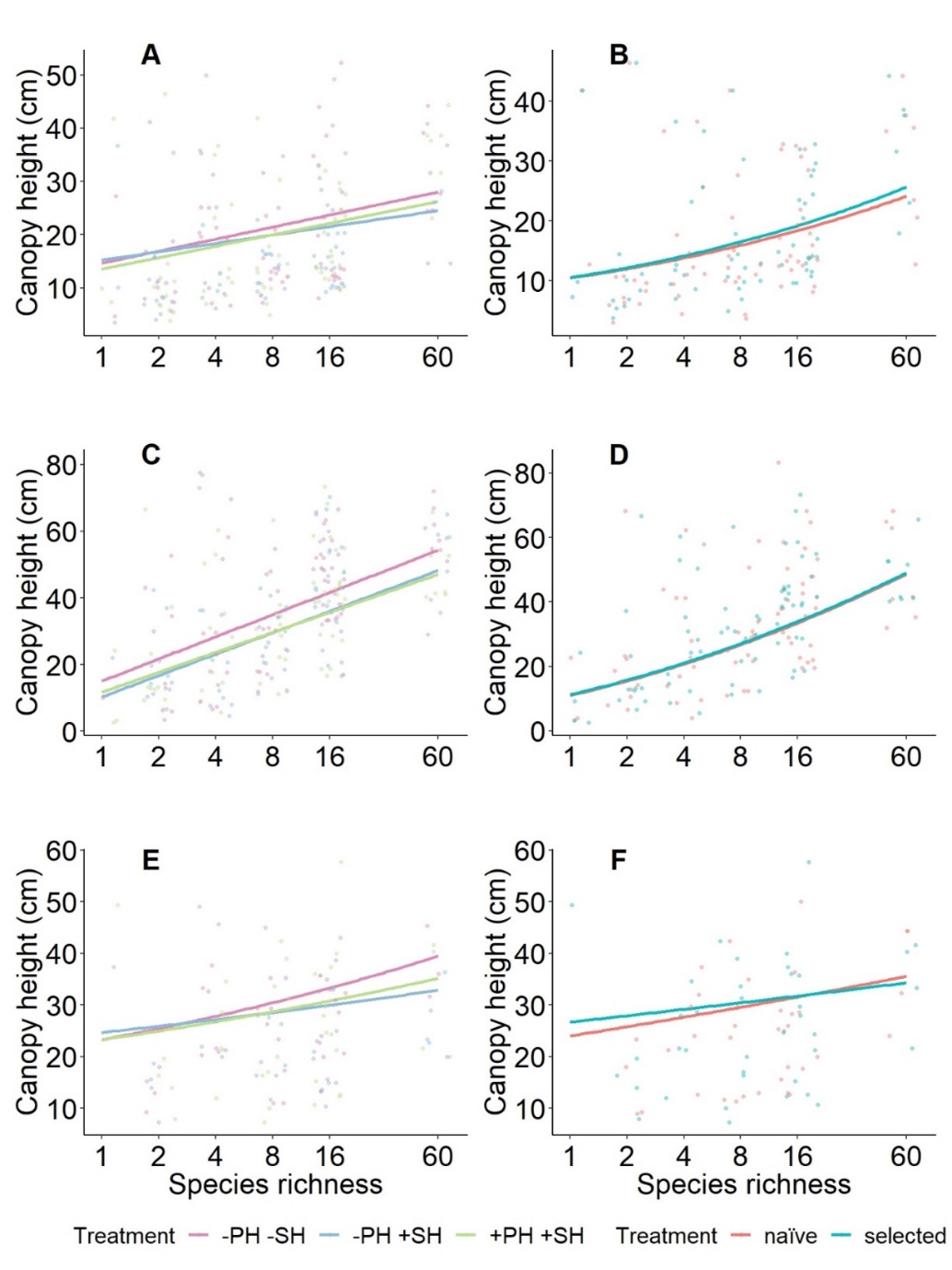
